# Blunted Fas signaling favors RIPK1-driven neutrophil necroptosis in critically ill COVID-19 patients

**DOI:** 10.1101/2021.03.19.436166

**Authors:** Tiziano A. Schweizer, Srikanth Mairpady Shambat, Clément Vulin, Sylvia Hoeller, Claudio Acevedo, Markus Huemer, Alejandro Gomez-Mejia, Chun-Chi Chang, Jeruscha Baum, Sanne Hertegonne, Eva Hitz, Daniel A. Hofmaenner, Philipp K. Buehler, Holger Moch, Reto A. Schuepbach, Silvio D. Brugger, Annelies S. Zinkernagel

## Abstract

Critically ill COVID-19 patients are characterized by a severely dysregulated cytokine profile and elevated neutrophil counts, which are thought to contribute to disease severity. However, to date it remains unclear how neutrophils contribute to pathophysiology during COVID-19. Here, we assessed the impact of the dysregulated cytokine profile on the tightly regulated cell death program of neutrophils. We show that in a subpopulation of neutrophils, canonical apoptosis was skewed towards rapidly occurring necroptosis. This phenotype was characterized by abrogated caspase-8 activity and increased RIPK1 levels, favoring execution of necroptosis via the RIPK1-RIPK3-MLKL axis, as further confirmed in COVID-19 biopsies. Moreover, reduction of sFas-L levels in COVID-19 patients and hence decreased signaling to Fas directly increased RIPK1 levels and correlated with disease severity. Our results suggest an important role for Fas signaling in the regulation of cell death program ambiguity via the ripoptosome in neutrophils during COVID-19 and a potential therapeutic target to curb inflammation and thus influence disease severity and outcome.

## Introduction

Patients experiencing the severe form of coronavirus disease 2019 (COVID-19), caused by SARS-CoV-2, are at elevated risk for succumbing to respiratory failure, which is linked to elevated mortality (Huang et al., 2020; Wunsch, 2020; Wendel Garcia et al., 2020). The current state of knowledge indicates a strongly dysregulated immune response during critical COVID-19, involving different forms of regulated cell death (RCD), which affects a broad range of cell types, including neutrophils (Adamo et al., 2020; Althaus et al., 2020; Nagashima et al., 2020; Varga et al., 2020; Rodrigues et al., 2020; Middleton et al., 2020; Karki et al., 2021).

Death of neutrophils is a tightly regulated process. They usually die via apoptosis and are removed by efferocytosis (Lawrence et al., 2020). Lately, neutrophils have been investigated mostly for their capacity to employ extracellular traps (NETs), chromatin-based webs spiked with nuclear and cytosolic components, to trap and eliminate pathogens (Branzk et al., 2014). However, NETs have also been linked to a variety of non-infectious diseases, responsible for tissue-injury (Nakazawa et al., 2018). Inconclusive evidence suggests that neutrophils might contribute to immunothrombosis and lung-injury in critical COVID-19 due to higher rates of NETs, as indicated by the presence of either extracellular DNA co-localized with myeloperoxidase (MPO), citrullinated histones or elastase in tracheal aspirates and histological lung sections (Middleton et al., 2020; Radermecker et al., 2020; Nathan, 2020; Protasio Veras et al., 2020). However, intracellular content might also be released during regulated necrosis, termed necroptosis (D’Cruz et al., 2018; Nakazawa et al., 2018). Necroptosis, as well as apoptosis, is initiated by ripoptosome assembly. The ripoptosome is a multiprotein signaling complex which can either induce apoptosis, necroptosis or promote cell survival, depending on its stoichiometric composition of the key components receptor interacting serine-threonine protein kinase (RIPK) 1, caspase-8, Fas associated protein with death domain (FADD) and cellular FADD-like IL-1β-converting enzyme inhibitory protein (cFLIP) isoforms (Feoktistova et al., 2011). In case of stabilized RIPK1, necroptosis is driven by RIPK1-RIPK3 necrosome formation and subsequent activation of the mixed lineage kinase like (MLKL) (Schilling et al., 2014), inducing cell rupture (Orozco et al., 2014; Wang et al., 2016). Assembly and functionality of the ripoptosome is strictly regulated, amongst others by the Fas/Fas-Ligand (Fas-L) system (Tummers et al., 2020). However, whether skewed RCD mechanisms of neutrophils contribute towards augmenting tissue injury and inflammation in COVID-19 is still not well understood. Importantly, given that neutrophil counts are significantly elevated in critically ill COVID-19 patients, necroptosis could possibly contribute to severe inflammation in these patients.

Here, we report that neutrophils undergo rapid necroptosis, due to increased RIPK1-dominant ripoptosome function and execution of necroptosis via RIPK3-MLKL, elicited by impaired Fas/soluble Fas-L (sFas-L) signaling in critically ill COVID-19 patients, which correlated with disease severity.

## Results and discussion

### Acute phase plasma from critically ill COVID-19 patients induces lytic regulated cell death in neutrophils

To investigate the fate of neutrophil cell death during COVID-19, we looked at critically ill patients enrolled in our prospective intensive care unit (ICU) cohort (**Table S1**). Neutrophils were isolated within the first four days upon ICU admission (acute phase). In parallel, neutrophils from healthy donors were isolated. Neutrophils were stimulated with auto- or heterologous plasma and assessed for short term viability using flow cytometry (**Fig. 1A and B, S1A**). Stimulation with acute phase COVID-19 plasma decreased the proportion of live cells and increased the proportion of RCD, as indicated by Annexin V positive staining, of both COVID-19 and healthy donor neutrophils. (**Fig. 1B and C**). Furthermore, pre-stimulation with COVID-19 plasma and subsequent bacterial challenge underscored the cell-death prone phenotype of both COVID-19 and healthy donor neutrophils in the COVID-19 environment, simulated by addition of COVID-19 plasma (**Fig. S1B**). In contrast, neutrophils from the same COVID-19 patients during recovery phase (discharged from ICU or SARS-CoV-2 negative in a non-critical state), or healthy donor neutrophils stimulated with recovery phase plasma, displayed no difference in proportion of viable and dying cells, as compared to stimulation with healthy plasma (**Fig. 1D**), highlighting a transient effect of acute phase COVID-19 plasma on RCD induction in neutrophils.

**Figure 1.**
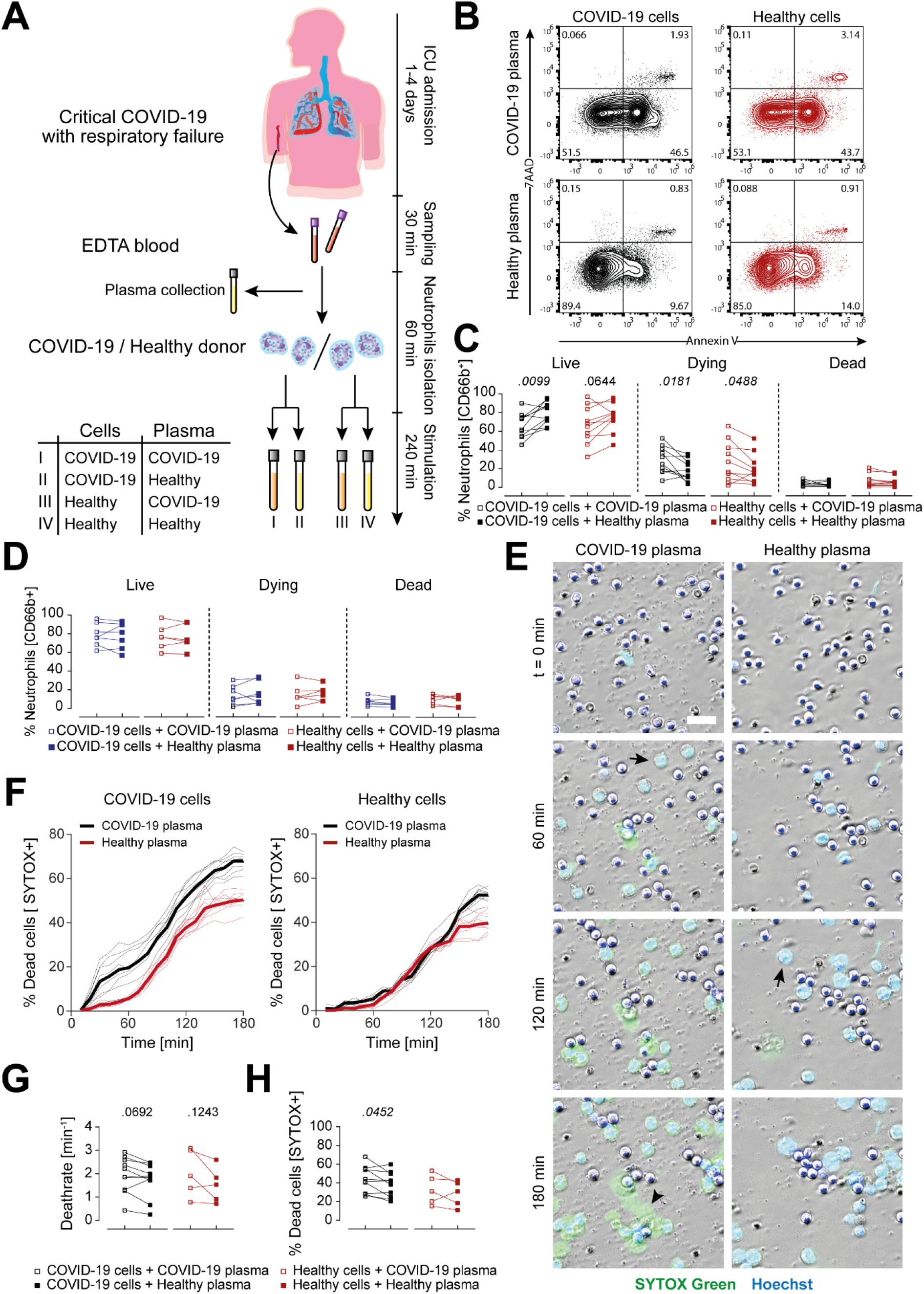
COVID-19 acute phase plasma induces lytic RCD of neutrophils. **(A)** Experimental overview. **(B and C)** Representative flow cytometry plots of acute COVID-19 (n=10) and healthy donor (n=10) neutrophils stimulated with auto- or heterologous plasma for 4 h (B) and quantification of live (Annexin V-/7AAD-), dying (Annexin V+/7AAD-) and dead (Annexin V+/7AAD+) neutrophils (C). **(D)** Quantification of live, dying and dead recovery COVID-19 (n=7) or healthy donor (n=6) neutrophils stimulated with auto- or heterologous plasma for 4 h. **(E)** Representative time lapse microscopy of acute COVID-19 neutrophils stimulated with auto- or heterologous plasma for 3 h. Cells were stained with Hoechst 33342 (blue) and SYTOX™ green. Images were taken every ten minutes. Scale bar, 30 μm. Arrows indicate cells with membrane breakdown, arrowheads indicate total cell lysis. **(F)** Representative cell death curve. Thin lines are FOV (n=8) per condition and thick line is mean of FOVs. Left: COVID-19 neutrophils, right: healthy neutrophils. **(G and H)** Quantification of cell death rate (G) and (H) proportion at 3 h of COVID-19 (n=10) or healthy neutrophils (n=5) stimulated with auto- or heterologous plasma. See also Videos 1 and 2. Connected squares represent one donor. Statistics were calculated by paired t-test or Wilcoxon signed-rank test. P values are indicated within the graphs. FOV, field of view.

To characterize the type of RCD the neutrophils succumbed to, we employed time lapse microscopy, using the cell-permeable DNA-dye Hoechst and the cell-impermeable DNA-dye SYTOX™ green. This approach revealed that COVID-19 plasma stimulation induced lytic RCD, linked to expulsion of DNA around the lysed cells, but distinct from classical NETs formation, while less cell lysis was observed upon stimulation with healthy plasma (**Fig. 1E, S1, C and D, Video 1 and 2**). This was in accordance with findings by Middleton et al., describing increased release of DNA from *ex vivo* cultured COVID-19 or healthy donor neutrophils stimulated with COVID-19 plasma, which they determined to be NETs (Middleton et al., 2020). Recently, we showed that neutrophils from COVID-19 patients released significantly fewer NETs upon bacterial challenge, the main natural inducer of NETs (Van Der Linden et al., 2017), questioning the presence of classical NETs in COVID-19 (Mairpady Shambat et al., 2020; Nathan, 2020). Protasio Veras et al. showed that SARS-CoV-2 can infect neutrophils, and viral replication caused release of NETs, thereby describing a situation different from cytokine-induced DNA release (Protasio Veras et al., 2020). The sensitivity of our method was further verified by neutrophils from a COVID-19 patient concomitantly treated with tamoxifen (**Fig. S1E**). Tamoxifen has been shown to augment the innate immune function of neutrophils as well as increase NETs formation (Corriden et al., 2015). In line with these previous findings, classical NETs (**Fig. S1F**) and apoptotic bodies were observed (**Fig. S1, G and H**). We analyzed cell death kinetics as proportion of SYTOX+ cells over time (**Fig. 1F and S2, A, B and C**) and verified that COVID-19 neutrophils died at a higher rate when stimulated with auto- as compared to heterologous plasma, resulting in a significantly increased proportion of dead cells (**Fig. 1, F, G and H**).

### COVID-19 plasma favors RIPK1-dominant ripoptosome phenotype

Since neutrophils usually die via caspase mediated apoptosis, we assessed whether caspases might be involved in the observed lytic RCD. Interestingly, caspase inhibition showed no effect on short term survival of neutrophils stimulated with COVID-19 plasma, confirming that neutrophils underwent caspase independent lytic RCD (**Fig. 2, A, B and S2D**). However, caspase blockage did not show any effect on short term survival upon healthy plasma stimulation, potentially due to the fact that caspases are usually only activated later on during the lifespan of neutrophils (Schwartz et al., 2012). Therefore, we also assessed long term survival in the COVID-19 and healthy plasma environment. Here, we found that COVID-19 plasma had rather the opposite outcome on long term as compared to short term stimulation, with higher proportions of neutrophils surviving as compared to healthy plasma (**Fig. 2, C and D**). Caspase inhibition was able to significantly elevate the proportion of surviving neutrophils in general. However, this effect was significantly lower when stimulated with COVID-19 plasma as compared to healthy plasma (**Fig. S2D**). This might be explained by elevated levels of G-CSF, GM-CSF and IL-8 during COVID-19 (Mairpady Shambat et al., 2020), which were described to interfere with caspase activation (van Raam et al., 2008; Klein et al., 2000). Additionally, certain viruses are known to secrete caspase-8 interfering peptides (Mocarski et al., 2012), which might also apply to SARS-CoV-2.

**Figure 2.**
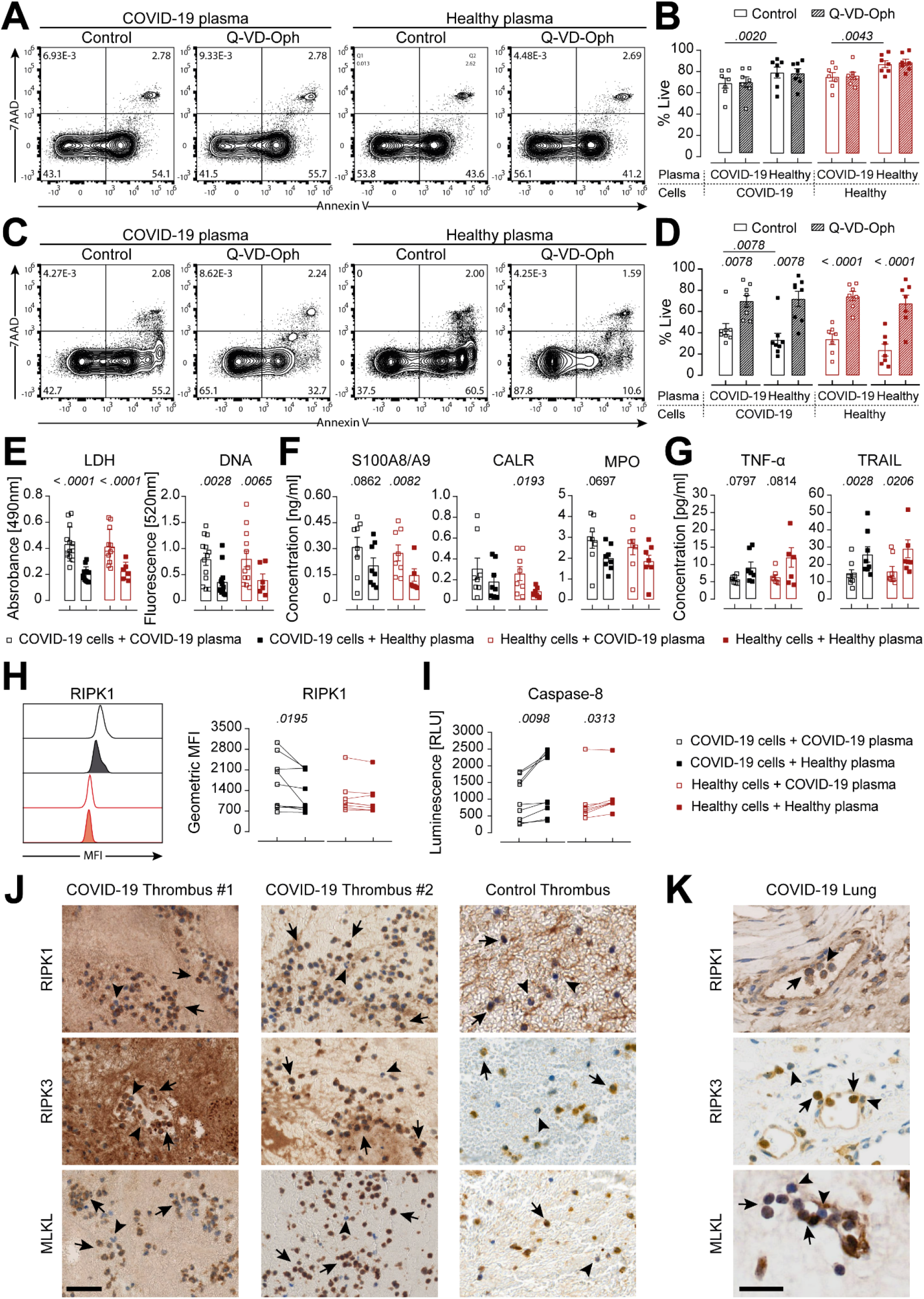
Neutrophil necroptosis via the RIPK1-RIPK3-MLKL axis. **(A-D)** Representative flow cytometry plots at 4 h (A) or 18 h post stimulation (C) and quantification of COVID-19 (n=7-8) or healthy donor (n=7) neutrophils stimulated with auto- or heterologous plasma, untreated or treated with Q-VD-Oph for 4 h (B) or 18 h (D). **(E-G)** Analysis of LDH and DNA (E) as well as DAMPs (F) and cytokines (G) in supernatants of COVID-19 (n=7-13) or healthy donor (n=6-7) neutrophils stimulated with auto- or heterologous plasma for 4 h. **(H and I)** Representative histogram and quantification of GMFI of intracellular RIPK1 expression and caspase-8 activity (I) of COVID-19 (n=9) or healthy donor (n=6-7) neutrophils stimulated with auto- or heterologous plasma for 4 h. Each square or connected squares represent one donor. Shown are mean ± SEM. Statistics were calculated by paired t-test or Wilcoxon signed-rank test. P values are indicated within the graphs. (**J and K**) RIPK1, RIPK3 and MLKL staining of COVID-19 (n = 2) and non-COVID-19 thrombi (J) as well as COVID-19 lung biopsies (K). Scale bars, 50 μm (J) and 25 μm (K). Arrows indicate strong positive staining, arrowheads indicate negative or weak positive staining. SEM, standard error of means.

A caspase-impaired environment might favor the occurrence of necroptosis over apoptosis. Necroptosis is described to differ from apoptotic cell death by release of damage associated molecular patterns (DAMPs) known to trigger inflammation and decreased release of classical cytokines, such as members of the TNF superfamily for neutrophils (Kearney and Martin, 2017). Therefore, we assessed the secretion of selected DAMPs and cytokines. COVID-19 plasma stimulation resulted in significantly higher lactate dehydrogenase (LDH), DNA, S100A8/A9 and calreticulin (CALR) release as well as slightly elevated MPO as compared to healthy plasma stimulation (**Fig. 2, E and F**). DAMPs, especially S100A8/A9, have previously been described to be important drivers of COVID-19 pathogenesis (Guo et al., 2021). Furthermore, decreased levels of TNF-α and TRAIL were detected (**Fig. 2G**), confirming a necroptotic secretion profile (Alvarez-Diaz et al., 2016; Newton et al., 2016). Of note, effectors of apoptosis and necroptosis are found within the ripoptosome, the decisive complex for activation of either pathway (Chen et al., 2018; Duprez et al., 2012; Feoktistova et al., 2011). In order to understand the modus operandi of the ripoptosome, we assessed intracellular RIPK1 levels as well as caspase-8 activity. Neutrophils derived directly from COVID-19 patients stimulated with autologous plasma displayed significantly higher RIPK1 levels (**Fig. 2H**) and at the same time decreased caspase-8 activity (**Fig. 2I**) as compared to healthy plasma stimulation. If either caspase-8 homodimerization or caspase-8 cFLIPL heterodimerization occurs, the resulting caspase-8 activity will be sufficient to degrade RIPK1 and prevent necrosome assembly (Schilling et al., 2014). However, increased RIPK1 levels and decreased caspase-8 activity upon COVID-19 plasma stimulation strongly suggested a RIPK1-dominant ripoptosome, favoring necrosome assembly and execution of necroptosis.

### Evidence for RIPK1-RIPK3-MLKL driven neutrophil necroptosis in COVID-19 thrombus and lung tissue

Critical COVID-19 is linked to endothelial damage and neutrophil-rich thrombi (Varga et al., 2020; Middleton et al., 2020). Therefore, we assessed whether neutrophil necroptosis occurred in COVID-19 thrombi by immunostaining for the necroptosis markers RIPK1, RIPK3 and MLKL (**Table S2**). Histological analysis within thrombus biopsies of two COVID-19 patients showed strong positive RIPK1, RIPK3 and MLKL staining for neutrophils (**Fig. 2J**), further corroborating the observed necroptotic *ex vivo* phenotype of neutrophils in the COVID-19 environment. We also observed evidence for neutrophil necroptosis in a thrombus from a non-COVID-19 patient (**Fig 2J**), in line with previous reports of activated platelets, driving neutrophil necroptosis (Nakazawa et al., 2018).

Furthermore, we assessed whether neutrophil necroptosis might also occur during the severe lung pathology observed in critical COVID-19. Indeed, we discovered strong positive staining of general necroptosis markers in lung biopsy tissue (**Fig. S3A**), and especially strong positive staining for neutrophils (**Fig. 2K**). Interestingly, neutrophils were always strongly stained for RIPK1, RIPK3 and MLKL within blood vessels (**Fig. S3B**), pointing towards contribution of neutrophil necroptosis within thrombi in the lung and its devastating effect during critical COVID-19 resulting in respiratory failure. Our histological findings are in line with studies showing that intracellular neutrophil content found in tissue can be released during necroptosis via the RIPK1-RIPK3-MLKL axis (Desai et al., 2016a). Of note, neutrophil necroptosis seems to be not uniquely attributable to COVID-19, as it has been described to occur during *S. aureus* pneumonia (Greenlee-Wacker et al., 2014; Zhou et al., 2018) and might be a phenomenon also influencing influenza A pneumonia (Zhu et al., 2018; Narasaraju et al., 2011), as well as in thrombus formation in general (Laridan et al., 2017), where NETs have been implied. There is accumulating evidence that DNA release after MLKL-mediated necroptosis and NETs formation via peptidylarginine deiminase 4 (PAD4) upon bacterial challenge differ drastically, especially in the light that neutrophils remain alive after NETs formation, but not after necroptosis (Branzk et al., 2014; Van Der Linden et al., 2017; Yipp et al., 2012). Controversially, MLKL was also reported to be able to activate PAD4 and cause NETs formation upon bacterial challenge (D’Cruz et al., 2018), highlighting the fact that DNA aggregates or NETs can be released via different mechanisms upon distinct stimuli, but might ultimately result in a similar outcome (Wang et al., 2018; Desai et al., 2016b). However, the interplay between the necroptotic cell death and NETs pathway remains to be elucidated.

### Abrogated Fas engagement favors high RIPK1 levels

Next, we sought to investigate the signaling modalities contributing towards RIPK1 mediated neutrophil necroptosis in COVID-19. Early on, COVID-19 has been characterized by a highly dysbalanced cytokine profile (Chevrier et al., 2021; Schulte-Schrepping et al., 2020), suggesting that the presence of host-derived mediators of ripoptosome function might also be altered. In our recently described COVID-19 ICU cohort, we detected significantly elevated TNF-α and IFN-α levels, both known to modulate RCD (Mairpady Shambat et al., 2020). However, the expression of the TNF RI and IFNAR1 receptor on neutrophils from COVID-19 patients and healthy donors showed no difference, minimizing their potential pivotal role (**Fig. S3C**).

Two other well-known death receptor pathways are Fas/Fas-L and TRAIL-R1/TRAIL. We found significantly lower sFas-L concentrations in plasma from COVID-19 patients as compared to healthy donors, which was in line with a recent study (Abers et al., 2021), and equal levels of TRAIL (**Fig. 3A**). Importantly, during the recovery phase of these patients, the concentration of sFas-L was significantly higher as compared to their acute phase, reaching levels of healthy donors (**Fig. S3D**). Assessing the corresponding receptors, Fas and TRAIL-R1, we observed significantly higher Fas expression on neutrophils derived from COVID-19 patients as compared to healthy donors, whereas TRAIL-R1 expression displayed no difference (**Fig. 3B**). It has only recently been described that Fas expression is also significantly elevated on T cells in COVID-19 (Bellesi et al., 2020; Filbin et al., 2020; Schultheiß et al., 2020; Zhu et al., 2020). The signaling modality of Fas-L to Fas during inflammation, but also homeostasis, is still not completely understood. Fas-L can occur as sFas-L but also as membrane-bound (mFas-L) (Herrero et al., 2011). sFas-L can either induce or prevent cell death, depending on various environmental factors, whereas mFas-L is solely thought to induce cell death (Hohlbaum et al., 2000; Nguyen and Russell, 2001; Suda et al., 1997; Tummers et al., 2020). Interestingly, the supernatant of COVID-19 neutrophils contained significantly lower concentrations of sFas-L when stimulated with auto- as compared to heterologous plasma (**Fig. 3C**). Flow cytometry analysis revealed that neutrophils stimulated with COVID-19 plasma produced significantly lower levels of Fas-L (**Fig. 3D**), confirming previous results that neutrophils express and secrete Fas-L (Serrao et al., 2001). It remains to be elucidated which other cell types apart from neutrophils contribute to the sFas-L pool found in steady state conditions and how their secretion profile is affected during COVID-19.

**Figure 3.**
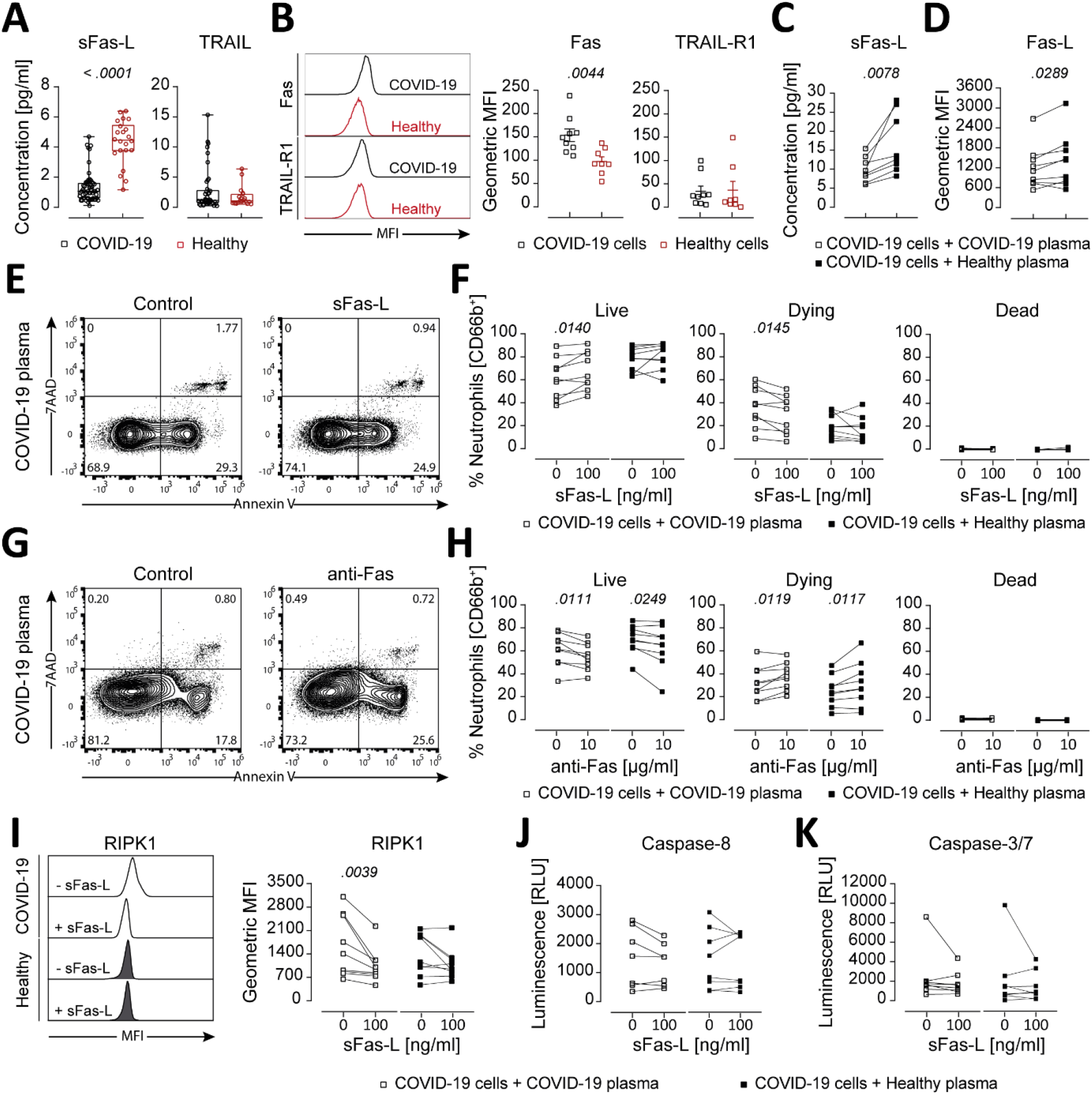
Impaired Fas signaling favors RIPK1-driven necroptosis. **(A)** Luminex-based analysis of COVID-19 (sFas-L, n=56/61; TRAIL, n=28/61 detected) and healthy donors’ plasma (sFas-L, n=22/22; TRAIL, n=17/22). **(B)** Representative histogram and quantification of receptor expression on neutrophils from COVID-19 (n=9) and healthy donors (n=8). **(C and D)** Luminex-based analysis of sFas-L in supernatants (C) and quantification of intracellular Fas-L expression (D) of COVID-19 neutrophils (n=8-9) stimulated with auto- or heterologous plasma for 4 h. **(E and F)** Representative flow cytometry plots (E) and quantification of live (Annexin V-/7AAD-), dying (Annexin V+/7AAD-) and dead (Annexin V+/7AAD+) COVID-19 neutrophils (n=9) stimulated with auto- or heterologous plasma and with or without 100 ng/ml sFas-L for 4 h (F). **(G and H)** Representative flow cytometry plots (G) and quantification of live, dying and dead COVID-19 neutrophils (n=9) stimulated with auto- or heterologous plasma and with or without 10 μg/ml anti-Fas for 4 h. **(I-K)** Representative histogram and quantification of GMFI of intracellular RIPK1 expression (I), caspase-8 activity (J) or caspase-3/7 activity (K) of COVID-19 neutrophils (n=8-9) stimulated with auto- or heterologous plasma and with or without 100 ng/ml sFas-L for 4 h. Each dot, square or connected squares represent one donor. Shown are mean ± SEM. Statistics were calculated by unpaired t-test, Mann-Whitney, paired t-test or Wilcoxon signed-rank test. P values are indicated within the graphs. SEM, standard error of means.

We reasoned that sFas-L might be beneficial for survival of COVID-19 neutrophils. Indeed, addition of recombinant human sFas-L to COVID-19 neutrophils significantly increased the proportion of live cells and decreased RCD induction in the COVID-19 environment (**Fig. 3, E and F**). Blocking sFas-L from binding to Fas by employing a Fas-blocking antibody on the other hand increased the rate of RCD significantly, also during healthy plasma stimulation, confirming that binding of sFas-L to Fas is required for enhanced survival of COVID-19 neutrophils (**Fig. 3, G and H**). Next, we assessed whether sFas-L stimulation influenced RIPK1 levels and caspase-8 activity. Indeed, we found that treatment with sFas-L decreased RIPK1 levels significantly during COVID-19, but not healthy, plasma stimulation (**Fig. 3I**). This occurred independently of caspase-8 (**Fig. 3J**) and caspase-3/7 activity (**Fig. 3K**), suggesting that sFas-L acted as a pro-survival signal in the COVID-19 environment.

### Low sFas-L levels correlate with disease severity

Finally, we evaluated whether sFas-L levels, and hence neutrophil necroptosis, correlated with clinical status of the corresponding patients. Analysis of key markers for COVID-19 severity, i.e. neutrophil to lymphocyte ratio (NLR), monocyte counts, IL-6, CRP, bilirubin, platelet counts, length of ventilation (LOV) and length of ICU stay (LOS) (Qin et al., 2020; Carissimo et al., 2020; Hazard et al., 2020) to sFas-L levels did not display any significant correlation (**Fig. S3, E-L**). Of note, on the 17^th^ of July 2020, the RECOVERY trial showed reduced 28-day mortality in ventilated COVID-19 patients treated with dexamethasone (Horby et al., 2020), which then became standard of care (SOC) also at the University Hospital Zurich’s (USZ) ICU. Since dexamethasone is known to impact neutrophil functions and lifespan (Ronchetti et al., 2018), we performed an in-depth analysis comparing patients treated at attending physicians’ discretion (USZ SOC) or RECOVERY SOC. We identified significant negative correlation between sFas-L levels and NLR (**Fig. 4A**) but not monocyte counts (**Fig. 4B**), as well as IL-6 (**Fig. 4C**), but not CRP (**Fig. 4D**) in the USZ SOC group. These findings might suggest that neutrophil necroptosis due to decreased sFas-L levels favors an inflammation feedback loop with a central role for IL-6 signaling, potentially sustaining emergency granulopoiesis and aborting lymphopoiesis, which would explain the increased NLR (Maeda et al., 2009). However, this remains speculative and further research is needed to clarify how decreased sFas-L impacts viability of lymphocytes. Furthermore, low sFas-L levels also correlated with high bilirubin levels (**Fig. 4E**) and low platelet counts (**Fig. 4F**), further highlighting the devastating effect of neutrophil necroptosis on coagulation and tissue damage, as has been shown to impact COVID-19 severity (Liu et al., 2020; Radermecker et al., 2020). In line with these findings, low sFas-L levels were linked to increased LOV (**Fig. 4G**) and also LOS (**Fig. 4H**) in the USZ SOC group. We observed no correlation of sFas-L to disease severity in the RECOVERY SOC, even though sFas-L levels were similar in RECOVERY as well as USZ SOC group (**Fig. S3M**). Synthetic glucocorticoids, such as dexamethasone, are known to favor neutrophil maturation and tissue retention (Ronchetti et al., 2018). Therefore, glucocorticoids might ameliorate disease severity by either accelerating neutrophil maturation or dampening the inflammatory COVID-19 environment, leaving them less prone for sFas-L dependent necroptosis. Indeed, when comparing the viability of neutrophils from COVID-19 patients treated under USZ or RECOVERY SOC (as shown in **Fig. 1C**), the later showed enhanced viability (**Fig. S3N**). However, prescription of synthetic glucocorticoids during viral pneumonia is still controversial, as, among other factors, the timing, dosage and duration of application play an important role defining the potential beneficial or detrimental effect on patient outcome (Yang et al., 2020).

**Figure 4.**
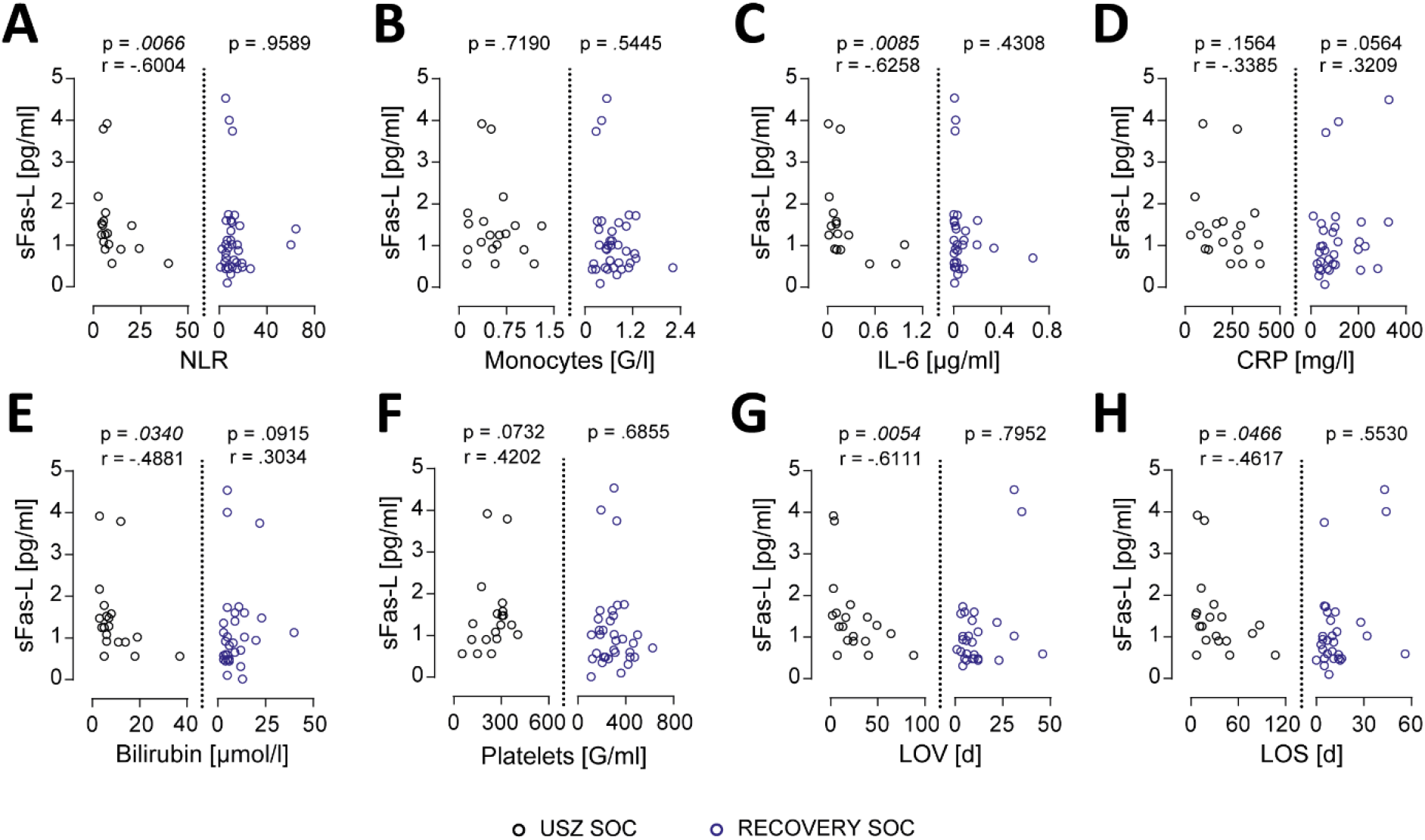
Correlation of low sFas-L levels with disease severity and abolishment by RECOVERY SOC. **(A-H)** Correlation analysis of plasma sFas-L levels with NLR (A), monocytes (B), IL-6 (C), CRP (D), bilirubin (E), platelets (F), LOV (G) and LOS (H) in the USZ (n=19) and RECOVERY SOC (n=36) group. Each dot represents one donor for which sFas-L was detected (n=55). One patient from the RECOVERY SOC group received dexamethasone only after experimental sampling and was excluded from correlation analysis. Statistics were calculated by non-parametric Spearman correlation.

Although the findings presented in this work were obtained from a small cohort at a single center only, they nevertheless further elucidate the crucial role of neutrophils during COVID-19 and deliver novel insights into the important regulation of the ripoptosome by the Fas/Fas-L system and its correlation to disease severity. Our findings provide hints for future potential therapeutic development, aiming at restoring the fate of neutrophils and benefiting patient outcome, potentially also beyond COVID-19.

## Materials and methods

### Patients

Patients were recruited between April and December 2020 in the MicrobiotaCOVID prospective cohort study conducted at the Institute of Intensive Care Medicine of the University Hospital Zurich (USZ, Zurich, Switzerland) and were included in an extended subcohort as described previously (Mairpady Shambat et al., 2020; Buehler et al., 2021). The study was approved by the local ethics committee of the Canton of Zurich, Switzerland (Kantonale Ethikkommission Zurich BASEC ID 2020 – 00646) and is registered at clinicaltrials.gov (ClinicalTrials.gov Identifier: NCT04410263). Patients were considered to be in the acute phase within the first four days upon initial ICU admission, the recovery phase was defined as patients being discharged from the ICU or negative for SARS-CoV-2 and in a non-critical state. Blood sampling was carried out with EDTA tubes. Patient demographics and clinical as well as laboratory parameters are listed in **Table S1**.

### Plasma collection

EDTA tubes were centrifuged at 1800 rpm for 10 min (no acceleration and brakes) after which the plasma was separated from the cellular fraction and collected in a fresh tube. The cellular fraction of the blood was used for neutrophil isolation as described below. The collected plasma was centrifuged again at 3000 rpm for 10 min (full acceleration and brakes) to pellet debris and the clear plasma supernatant was collected. Plasma samples were either used directly for the experiments or aliquoted and frozen at -80°C for cytokine analysis.

### Cytokine analysis

Cytokine levels in plasma from COVID-19 patients and healthy donors, as well as cell culture supernatants were analyzed on a Luminex™ MAGPIX™ instrument with a custom human cytokine panel (ThermoFisher). Samples were thawed at room temperature and prepared according to the manufacturer’s instructions. In brief, magnetic beads were added to the 96-well plate on a magnetic holder and incubated for 2 min. The plate was washed twice with assay buffer for 30 sec each. Provided standards were diluted in assay buffer or RPMI (Gibco) for analysis of plasma levels or cell culture supernatants, respectively. Cell culture supernatants were measured undiluted, whereas plasma samples were diluted 1:2 in assay buffer. The plate was incubated for 2 h at room temperature (RT) at 550 rpm in an orbital plate shaker. Next, the plate was washed twice and incubated for 30 min at 550 rpm with detection antibodies. Following further washing steps, the plate was incubated with Streptavidin-PE solution for 30 min at 550 rpm. Finally, the plate was washed, reading buffer was added and incubated for 10 min at RT and 550 rpm before running the plate. Analysis was performed using the xPONENT® software. Data was validated additionally with the ProcartaPlex Analyst software (ThermoFisher).

### Neutrophils isolation and plasma stimulation

Neutrophils from COVID-19 patients and healthy donors were isolated with the EasySep™ Direct Human Neutrophil Isolation Kit (StemCell™) according to the manufacturer’s instructions. In brief, the cellular fraction of the blood was diluted 1:2 with Dulbecco’s phosphate buffered saline (DPBS, Gibco) and neutrophil enrichment cocktail was added for 15 min. Next, the magnetic beads were added for 15 min. The samples were once more diluted 1:2 with DPBS and placed in a magnetic holder (StemCell) for 15 min. Neutrophils were collected and centrifuged for 1500 rpm (low acceleration and brakes) for 5 min. Red blood cells lysis was performed with H2O and stopped with DPBS after which the samples were centrifuged. Neutrophils were resuspended in RPMI and counted on an Attune NxT (ThermoFisher). Neutrophils were either directly stained for cell surface receptor analysis using flow cytometry or prepared in RPMI containing 10% either auto- or heterologous plasma and seeded in either V-well canonical plates (Corning, for flow cytometry analysis) or 8-well microslides (ibidi, for microscopy time lapse). Plates were incubated for 4 h-18 h at 37°C + 5% CO2. For bacterial challenge, plates were incubated for 2.5 h after which neutrophils were challenged with opsonized (plasma-specific, 20 min at RT) bacteria at MOI 10 for 1.5 h.

### Bacterial strains

The *Staphylococcus aureus* strain JE2 (MRSA-USA300) (Frey et al., 2021) was grown in Tryptic Soy Broth at 37°C and 220 rpm for 16 h. Cultures were diluted in fresh TSB and grown to exponential phase for the challenge. *Streptococcus pneumoniae* (serotype 6B) (Malley et al., 2001) was passaged twice on blood agar plates (Columbia blood agar, Biomereux) and incubated at 37°C with 5% CO2. A liquid culture was prepared in Todd Hewitt Yeast broth with a starting OD600nm of 0.1 and grown at 37°C in a water bath until OD600nm of 0.35 for the challenge.

### Cell death analysis by flow cytometry

If indicated, pan-caspase inhibitor Q-VD-Oph (50 μM, Sigma-Aldrich), recombinant human sFas-L (100 ng/ml, Enzo Lifescience), Ultra-LEAF™ purified anti-human CD95 (FAS) blocking antibody (10 μg/ml, clone A16086F, Biolegend) or respective DMSO and H2O control were added prior to incubation. After the incubation period, plates were centrifuged at 1500 rpm (full acceleration and brakes) for 6 min. Supernatants were collected for further analyses and the wells were washed once with FACS buffer (DPBS + 5% fetal calf serum and 1 mM EDTA). Cells were stained with anti-CD66b APC (G10F5) from Thermofisher in FACS buffer for 30 min at 4°C. The wells were washed once with Annexin V buffer (Biolegend) and cells were stained with Annexin V FITC and 7AAD (Biolegend) for 30 min at RT. The plates were acquired on an Attune NxT. The gating strategy is depicted in **Fig. S1A**.

### Lactate dehydrogenase release measurement

Supernatants were incubated 1:2 with the substrate solution of the CytoTox 96® Non-Radioactive Cytotoxicity Assay (Promega) for 30 min in the dark in a 96-well plate (Greiner), after which the stop solution was added. The resulting absorbance of the converted substrate due to released lactate dehydrogenase (LDH) was measured at 490 nm.

### DNA-release measurement

Supernatants were incubated 1:2 with 60nM SYTOX™ Green (ThermoFisher) for 30min at 4°C in a 96-well black bottom plate (Greiner). The fluorescence of the bound DNA was measured in a fluorescence plate reader (Molecular Probes) with excitation at 488 nm and emission at 520 nm.

### Cell surface receptors and ligand analysis by flow cytometry

To assess intracellular Fas-L levels, GolgiStop™ (BD Biosciences) was added according to the manufacturer’s instruction for 4 h. After the incubation period, plates were centrifuged and washed once with DPBS. Cells were stained with the Fixable LIVE/DEAD™ Near-IR dead cell marker (ThermoFisher) in DPBS for 25 min at 4°C. Next, the wells were washed once with FACS buffer and cells were stained with anti-CD66b APC or PE-Cy7 (G10F5) and anti-IFNAR-1 PE (MAR1-5A3) from ThermoFisher, anti-Fas BV421 (DX2), anti-TRAIL-R1 APC (DJR1), anti-TNF RI PE (W15099A) all from Biolegend in FACS buffer for 30 min at 4°C. For intracellular staining, the cells were fixed for 15 min at 4°C in the Fix/Perm Solution A from Fix/perm Kit. Next, staining with anti-Fas-L BV421 (NOK-1) from Biolegend or anti-RIPK1 AF488 (Polyclonal) from Bioss Antibodies was carried out in Fix/Perm solution B for 30 min at 4°C. The plates were acquired on an Attune NxT.

### Microscopy time lapse

After adding the cells to the microslides, 150 nM SYTOX™ Green (ThermoFisher) and 2 μM Hoechst 33342 (ThermoFisher) were added and the slides were centrifuged at 1500 rpm (low acceleration and brakes) for 3 min. Microslides were imaged for 4 h at 37°C on a fully automated Olympus IX83 microscope with a 40X objective (UPLFLN40XPH-2), illuminated with a PE-4000 LED system through a quadband filter set (U-IFCBL50). Eight observation positions per well (condition) were assigned before the time lapse was started to avoid potential observer bias. The proportion of dead neutrophils was assessed as described in **Fig. S2A**: after filtering the nuclei on the Hoechst signal and watershed segmentation, cells were tracked through time and either assigned the category “live” or “dead” based on the value of the SYTOX Green signal. Assigned dead cells were counted until the end of the experiment, even after the disappearance of the fluorescent signal. The exponential death rate was fitted from the first few hours of the experiment. Images were processed using ImageJ software (Schneider et al., 2012).

### Histology

Formalin-fixed paraffin-embedded tissue sections were pre-treated with Tris-EDTA buffer (pH9.0) at 100°C for 30min. Next, they were stained with RIPK1 (ab72139, clone 7H10, dilution 1:1500), RIPK3 (ab62344, polyclonal, dilution 1:100) and MLKL (ab184718, EPR17514, dilution 1:100), all from abcam for 30 min. For detection, the slides were stained with the Bond Refine Detection Kit (Leica Biosystems), according to the manufacturer’s instruction. Finally, the slides were counterstained with haematoxylin. For background information of the obtained biopsies, see **Table S2**. The non-COVID-19 thrombus section was obtained from a patient with a general consent.

### Statistical analyses

The number of donors can be found in the corresponding figure legends. Samples were assessed for normal distribution. Differences between two groups were calculated using either unpaired t-test, Mann-Whitney, paired t-test or Wilcoxon signed-rank test in Prism (GraphPad). Correlation of clinical parameters was computed using non-parametric Spearman correlation in Prism.

## Acknowledgments

We’d like to thank our patients for willing to participate in this study. Furthermore, we’d like to thank André Fitsche, Christiane Mittmann and Susanne Dettwiler for their help with histology.

## Funding

This work was funded by the SNSF project grant 31003A_176252 (to A.S.Z), the SNF Biobanking grant 31BK30_185401 (to A.S.Z), the Uniscientia Foundation Grant (to A.S.Z), the Swedish Society for Medical Research (SSMF) foundation grant P17-0179 (to S.M.S), the Promedica Foundation 1449/M (to S.D.B) and unrestricted funds (to R.A.S).

## Author contributions

TA. Schweizer, S. Mairpady Shambat and AS. Zinkernagel conceived the project and were involved in experimental design. TA. Schweizer performed most experiments, analyzed and compiled the data with help of S. Mairpady Shambat. C. Vulin performed microscopy and analysis. M. Huemer, A. Gomez-Mejia, CC. Chang, J. Baum and S. Hertegonne assisted in experiments and specimen processing. TA. Schweizer prepared the figures with help of C. Vulin. S. Hoeller and H. Moch performed histology. C. Acevedo, E. Hitz, DA. Hofmaenner, PK. Buehler and SD. Brugger carried out patient recruitment, collection of patient specimens and epidemiological as well as clinical data. TA. Schweizer and S. Mairpady Shambat wrote the first draft of the manuscript. All authors helped in editing the final version of the manuscript and approved it.

## Disclosures

The authors declare no competing financial interests.

## Notes

### Competing Interest Statement

The authors have declared no competing interest.

## References

Abers, M.S., O.M. Delmonte, E.E. Ricotta, J. Fintzi, D.L. Fink, A.A. Almeida de Jesus, K.A. Zarember, S. Alehashemi, V. Oikonomou, J. V. Desai, S.W. Canna, B. Shakoory, K. Dobbs, L. Imberti, A. Sottini, E. Quiros-Roldan, F. Castelli, C. Rossi, D. Brugnoni, A. Biondi, L.R. Bettini, M. D’Angio’, P. Bonfanti, R. Castagnoli, D. Montagna, A. Licari, G.L. Marseglia, E.F. Gliniewicz, E. Shaw, D.E. Kahle, A.T. Rastegar, M. Stack, K. Myint-Hpu, S.L. Levinson, M.J. DiNubile, D.W. Chertow, P.D. Burbelo, J.I. Cohen, K.R. Calvo, J.S. Tsang, H.C. Su, J.I. Gallin, D.B. Kuhns, R. Goldbach-Mansky, M.S. Lionakis, and L.D. Notarangelo. 2021. An immune-based biomarker signature is associated with mortality in COVID-19 patients. JCI Insight. 6. doi:10.1172/jci.insight.144455.

Adamo, S., S. Chevrier, C. Cervia, Y. Zurbuchen, M.E. Raeber, L. Yang, S. Sivapatham, A. Jacobs, E. Bächli, A. Rudiger, M. Stüssi-Helbling, L.C. Huber, D.J. Schaer, B. Bodenmiller, O. Boyman, and J. Nilsson. 2020. Lymphopenia-induced T cell proliferation is a hallmark of severe COVID-19. bioRxiv. doi:10.1101/2020.08.04.236521.

Althaus, K., I. Marini, J. Zlamal, L. Pelzl, A. Singh, H. Hä berle, M. Mehrlä nder, S. Hammer, H. Schulze, M. Bitzer, N. Malek, D. Rath, H. Bösmüller, B. Nieswandt, M. Gawaz, T. Bakchoul, and P. Rosenberger. 2020. Antibody-induced procoagulant platelets in severe COVID-19 infection. Blood. doi:10.1182/blood.2020008762.

Alvarez-Diaz, S., C.P. Dillon, N. Lalaoui, M.C. Tanzer, D.A. Rodriguez, A. Lin, M. Lebois, R. Hakem, E.C. Josefsson, L.A. O’Reilly, J. Silke, W.S. Alexander, D.R. Green, and A. Strasser. 2016. The Pseudokinase MLKL and the Kinase RIPK3 Have Distinct Roles in Autoimmune Disease Caused by Loss of Death-Receptor-Induced Apoptosis. Immunity. 45:513–526. doi:10.1016/j.immuni.2016.07.016.

Bellesi, S., E. Metafuni, S. Hohaus, E. Maiolo, F. Marchionni, S. D’Innocenzo, M. La Sorda, M. Ferraironi, F. Ramundo, M. Fantoni, R. Murri, A. Cingolani, S. Sica, A. Gasbarrini, M. Sanguinetti, P. Chiusolo, and V. De Stefano. 2020. Increased CD95 (Fas) and PD-1 expression in peripheral blood T lymphocytes in COVID-19 patients. Br. J. Haematol. 95:1–5. doi:10.1111/bjh.17034.

Branzk, N., A. Lubojemska, S.E. Hardison, Q. Wang, M.G. Gutierrez, G.D. Brown, and V. Papayannopoulos. 2014. Neutrophils sense microbe size and selectively release neutrophil extracellular traps in response to large pathogens. Nat. Immunol. 15:1017–25. doi:10.1038/ni.2987.

Buehler, P.K., A.S. Zinkernagel, D.A. Hofmaenner, P. David, W. Garcia, C.T. Acevedo, A. Gómez-mejia, S. Mairpady, F. Andreoni, M.A. Maibach, J. Bartussek, M.P. Hilty, P.M. Frey, R.A. Schuepbach, and S.D. Brugger. 2021. Bacterial pulmonary superinfections are associated with unfavourable outcomes in critically ill COVID-19 patients. Cell Reports Med. 100229. doi:10.1016/j.xcrm.2021.100229.

Carissimo, G., W. Xu, I. Kwok, M.Y. Abdad, Y.H. Chan, S.W. Fong, K.J. Puan, C.Y.P. Lee, N.K.W. Yeo, S.N. Amrun, R.S.L. Chee, W. How, S. Chan, B.E. Fan, A.K. Andiappan, B. Lee, O. Rötzschke, B.E. Young, Y.S. Leo, D.C. Lye, L. Renia, L.G. Ng, A. Larbi, and L.F. Ng. 2020. Whole blood immunophenotyping uncovers immature neutrophil-to-VD2 T-cell ratio as an early marker for severe COVID-19. Nat. Commun. 11:1–12. doi:10.1038/s41467-020-19080-6.

Chen, K.W., K.E. Lawlor, J.B. von Pein, D. Boucher, M. Gerlic, B.A. Croker, J.S. Bezbradica, J.E. Vince, and K. Schroder. 2018. Cutting Edge: Blockade of Inhibitor of Apoptosis Proteins Sensitizes Neutrophils to TNF-but Not Lipopolysaccharide-Mediated Cell Death and IL-1β Secretion. J. Immunol. 200:3341–3346. doi:10.4049/jimmunol.1701620.

Chevrier, S., Y. Zurbuchen, C. Cervia, S. Adamo, M.E. Raeber, N. de Souza, S. Sivapatham, A. Jacobs, E. Bachli, A. Rudiger, M. Stüssi-Helbling, L.C. Huber, D.J. Schaer, J. Nilsson, O. Boyman, and B. Bodenmiller. 2021. A distinct innate immune signature marks progression from mild to severe COVID-19. Cell Reports Med. 2. doi:10.1016/j.xcrm.2020.100166.

Corriden, R., A. Hollands, J. Olson, J. Derieux, J. Lopez, J.T. Chang, D.J. Gonzalez, and V. Nizet. 2015. Tamoxifen augments the innate immune function of neutrophils through modulation of intracellular ceramide. Nat. Commun. 6:8369. doi:10.1038/ncomms9369.

D’Cruz, A.A., M. Speir, M. Bliss-Moreau, S. Dietrich, S. Wang, A.A. Chen, M. Gavillet, A. Al-Obeidi, K.E. Lawlor, J.E. Vince, M.A. Kelliher, R. Hakem, M. Pasparakis, D.A. Williams, M. Ericsson, and B.A. Croker. 2018. The pseudokinase MLKL activates PAD4-dependent NET formation in necroptotic neutrophils. Sci. Signal. 11:1–12. doi:10.1126/scisignal.aao1716.

Desai, J., S. V. Kumar, S.R. Mulay, L. Konrad, S. Romoli, C. Schauer, M. Herrmann, R. Bilyy, S. Müller, B. Popper, D. Nakazawa, M. Weidenbusch, D. Thomasova, S. Krautwald, A. Linkermann, and H.J. Anders. 2016a. PMA and crystal-induced neutrophil extracellular trap formation involves RIPK1-RIPK3-MLKL signaling. Eur. J. Immunol. 46:223–229. doi:10.1002/eji.201545605.

Desai, J., S.R. Mulay, D. Nakazawa, and H.J. Anders. 2016b. Matters of life and death. How neutrophils die or survive along NET release and is “NETosis” = necroptosis? Cell. Mol. Life Sci. 73:2211–2219. doi:10.1007/s00018-016-2195-0.

Duprez, L., M.J.M. Bertrand, T. Vanden Berghe, Y. Dondelinger, N. Festjens, and P. Vandenabeele. 2012. Intermediate domain of Receptor-interacting Protein Kinase 1 (RIPK1) determines switch between necroptosis and RIPK1 kinase-dependent apoptosis. J. Biol. Chem. 287:14863–14872. doi:10.1074/jbc.M111.288670.

Feoktistova, M., P. Geserick, B. Kellert, D.P. Dimitrova, C. Langlais, M. Hupe, K. Cain, M. MacFarlane, G. Häcker, and M. Leverkus. 2011. CIAPs Block Ripoptosome Formation, a RIP1/Caspase-8 Containing Intracellular Cell Death Complex Differentially Regulated by cFLIP Isoforms. Mol. Cell. 43:449–463. doi:10.1016/j.molcel.2011.06.011.

Filbin, M.R., A. Mehta, A.M. Schneider, K.R. Kays, and J.R. Guess. 2020. Plasma proteomics reveals tissue-specific cell death and mediators of cell-cell interactions in severe COVID-19 patients. bioRxiv. doi:10.1101/2020.11.02.365536.

Frey, P.M., J. Baer, J. Bergada-Pijuan, C. Lawless, P.K. Bühler, R.D. Kouyos, K.P. Lemon, A.S. Zinkernagel, and S.D. Brugger. 2021. Quantifying Variation in Bacterial Reproductive Fitness: a High-Throughput Method. mSystems. 6:1–13. doi:10.1128/msystems.01323-20.

Greenlee-Wacker, M.C., K.M. Rigby, S.D. Kobayashi, A.R. Porter, F.R. DeLeo, and W.M. Nauseef. 2014. Phagocytosis of Staphylococcus aureus by Human Neutrophils Prevents Macrophage Efferocytosis and Induces Programmed Necrosis. J. Immunol. 192:4709– 4717. doi:10.4049/jimmunol.1302692.

Guo, Q., Y. Zhao, J. Li, J. Liu, X. Yang, X. Guo, M. Kuang, H. Xia, Z. Zhang, L. Cao, Y. Luo, L. Bao, X. Wang, X. Wei, W. Deng, N. Wang, L. Chen, J. Chen, H. Zhu, R. Gao, C. Qin, X. Wang, and F. You. 2021. Induction of alarmin S100A8/A9 mediates activation of aberrant neutrophils in the pathogenesis of COVID-19. Cell Host Microbe. 29:222-235.e4. doi:10.1016/j.chom.2020.12.016.

Hazard, D., K. Kaier, M. Von Cube, M. Grodd, L. Bugiera, J. Lambert, and M. Wolkewitz. 2020. Joint analysis of duration of ventilation, length of intensive care, and mortality of COVID-19 patients: A multistate approach. BMC Med. Res. Methodol. 20:1–9. doi:10.1186/s12874-020-01082-z.

Herrero, R., X. Fu, T.R. Martin, R. Herrero, O. Kajikawa, G. Matute-bello, Y. Wang, N. Hagimoto, H. Rosen, R.B. Goodman, X. Fu, and T.R. Martin. 2011. The biological activity of FasL in human and mouse lungs is determined by the structure of its stalk region. J Clin Invest. 121:1174–1190. doi:10.1172/JCI43004.1174.

Hohlbaum, A.M., S. Moe, and A. Marshak-Rothstein. 2000. Opposing effects of transmembrane and soluble Fas ligand expression on inflammation and tumor cell survival. J. Exp. Med. 191:1209–1219. doi:10.1084/jem.191.7.1209.

Horby, P., L. Wei Shen, J.R. Emberson, M. Mafham, J.L. Bell, L. Linsell, N. Staplin, C. Brightling, A. Ustianowski, E. Elmahi, B. Prudon, C. Green, T. Felton, D. Chadwick, K. Rege, C. Fegan, L.C. Chappell, S.N. Faust, T. Jaki, K. Jeffery, A. Montgomery, K. Rowan, E. Juszczak, J.K. Baillie, R. Haynes, and M.J. Landray. 2020. Dexamethasone in Hospitalized Patients with Covid-19 — Preliminary Report. N. Engl. J. Med. 1–11. doi:10.1056/nejmoa2021436.

Huang, C., Y. Wang, X. Li, L. Ren, J. Zhao, Y. Hu, L. Zhang, G. Fan, J. Xu, X. Gu, Z. Cheng, T. Yu, J. Xia, Y. Wei, W. Wu, X. Xie, W. Yin, H. Li, M. Liu, Y. Xiao, H. Gao, L. Guo, J. Xie, G. Wang, R. Jiang, Z. Gao, Q. Jin, J. Wang, and B. Cao. 2020. Clinical features of patients infected with 2019 novel coronavirus in Wuhan, China. Lancet. 395:497–506. doi:10.1016/S0140-6736(20)30183-5.

Karki, R., B.R. Sharma, S. Tuladhar, E.P. Williams, L. Zalduondo, P. Samir, M. Zheng, B. Sundaram, B. Banoth, R.K.S. Malireddi, P. Schreiner, G. Neale, P. Vogel, R. Webby, C.B. Jonsson, and T.D. Kanneganti. 2021. Synergism of TNF-α and IFN-γ Triggers Inflammatory Cell Death, Tissue Damage, and Mortality in SARS-CoV-2 Infection and Cytokine Shock Syndromes. Cell. 184:149-168.e17. doi:10.1016/j.cell.2020.11.025.

Kearney, C.J., and S.J. Martin. 2017. An Inflammatory Perspective on Necroptosis. Mol. Cell. 65:965–973. doi:10.1016/j.molcel.2017.02.024.

Klein, J.B., M.J. Rane, J.A. Scherzer, P.Y. Coxon, R. Kettritz, J.M. Mathiesen, A. Buridi, and K.R. McLeish. 2000. Granulocyte-Macrophage Colony-Stimulating Factor Delays Neutrophil Constitutive Apoptosis Through Phosphoinositide 3-Kinase and Extracellular Signal-Regulated Kinase Pathways. J. Immunol. 164:4286–4291. doi:10.4049/jimmunol.164.8.4286.

Laridan, E., F. Denorme, L. Desender, O. François, T. Andersson, H. Deckmyn, K. Vanhoorelbeke, and S.F. De Meyer. 2017. Neutrophil extracellular traps in ischemic stroke thrombi. Ann. Neurol. 82:223–232. doi:10.1002/ana.24993.

Lawrence, S.M., R. Corriden, and V. Nizet. 2020. How Neutrophils Meet Their End. Trends Immunol. 41:531–544. doi:10.1016/j.it.2020.03.008.

Van Der Linden, M., G.H.A. Westerlaken, M. Van Der Vlist, J. Van Montfrans, and L. Meyaard. 2017. Differential Signalling and Kinetics of Neutrophil Extracellular Trap Release Revealed by Quantitative Live Imaging. Sci. Rep. 7:1–11. doi:10.1038/s41598-017-06901-w.

Liu, Z., J. Li, W. Long, W. Zeng, R. Gao, G. Zeng, D. Chen, S. Wang, Q. Li, D. Hu, L. Guo, Z. Li, and X. Wu. 2020. Bilirubin Levels as Potential Indicators of Disease Severity in Coronavirus Disease Patients: A Retrospective Cohort Study. Front. Med. 7:1–9. doi:10.3389/fmed.2020.598870.

Maeda, K., A. Malykhin, B.N. Teague-Weber, X.H. Sun, A.D. Farris, and K.M. Coggeshall. 2009. Interleukin-6 aborts lymphopoiesis and elevates production of myeloid cells in systemic lupus erythematosus-prone B6.Sle1.Yaa animals. Blood. 113:4534–4540. doi:10.1182/blood-2008-12-192559.

Mairpady Shambat, S., A. Gomez-Mejia, T.A. Schweizer, M. Huemer, C. Chang, C. Acevedo, J. Pijuan Bergada, C. Vulin, N. Miroshnikova, D.A. Hofmänner, P.D. Wendel Garcia, M.P. Hilty, R.A. Schüpbach, P.K. Bühler, S.D. Brugger, and A.S. Zinkernagel. 2020. Neutrophil and monocyte dysfunctional effector response towards bacterial challenge in critically-ill COVID-19 patients. bioRxiv. doi:doi.org/10.1101/2020.12.01.406306.

Malley, R., M. Lipsitch, A. Stack, R. Saladino, G. Fleisher, S. Pelton, C. Thompson, D. Briles, and P. Anderson. 2001. Intranasal immunization with killed unencapsulated whole cells prevents colonization and invasive disease by capsulated pneumococci. Infect. Immun. 69:4870–4873. doi:10.1128/IAI.69.8.4870-4873.2001.

Middleton, E.A., X. He, F. Denorme, R.A. Campbell, D. Ng, S.P. Salvatore, M. Mostyka, A. Baxter-stoltzfus, A.C. Borczuk, M. Loda, M.J. Cody, B.K. Manne, I. Portier, E.S. Harris, A.C. Petrey, E.J. Beswick, A.F. Caulin, A. Iovino, L.M. Abegglen, A.S. Weyrich, M.T. Rondina, M. Egeblad, J.D. Schiffman, C.C. Yost, and M. Elisa. 2020. Neutrophil extracellular traps contribute to immunothrombosis in COVID-19 acute respiratory distress syndrome. Blood. 136. doi:10.1182/blood.2020007008.

Mocarski, E.S., J.W. Upton, and W.J. Kaiser. 2012. Viral infection and the evolution of caspase 8-regulated apoptotic and necrotic death pathways. Nat. Rev. Immunol. 12:79–88. doi:10.1038/nri3131.

Nagashima, S., M.C. Mendes, A.P. Camargo Martins, N.H. Borges, T.M. Godoy, A.F.R.D.S. Miggiolaro, F. Da Silva Dezidério, C. Machado-Souza, and L. De Noronha. 2020. Endothelial dysfunction and thrombosis in patients with COVID-19 - Brief report. Arterioscler. Thromb. Vasc. Biol. 40:2404–2407. doi:10.1161/ATVBAHA.120.314860.

Nakazawa, D., J. Desai, S. Steiger, S. Müller, S.K. Devarapu, S.R. Mulay, T. Iwakura, and H.J. Anders. 2018. Activated platelets induce MLKL-driven neutrophil necroptosis and release of neutrophil extracellular traps in venous thrombosis. Cell Death Discov. 4. doi:10.1038/s41420-018-0073-2.

Narasaraju, T., E. Yang, R.P. Samy, H.H. Ng, W.P. Poh, A.A. Liew, M.C. Phoon, N. Van Rooijen, and V.T. Chow. 2011. Excessive neutrophils and neutrophil extracellular traps contribute to acute lung injury of influenza pneumonitis. Am. J. Pathol. 179:199–210. doi:10.1016/j.ajpath.2011.03.013.

Nathan, C. 2020. Neutrophils and COVID-19: Nots, NETs, and knots. J. Exp. Med. 217:3–5. doi:10.1084/jem.20201439.

Newton, K., D.L. Dugger, A. Maltzman, J.M. Greve, M. Hedehus, B. Martin-Mcnulty, R.A.D. Carano, T.C. Cao, N. Van Bruggen, L. Bernstein, W.P. Lee, X. Wu, J. Devoss, J. Zhang, S. Jeet, I. Peng, B.S. McKenzie, M. Roose-Girma, P. Caplazi, L. Diehl, J.D. Webster, and D. Vucic. 2016. RIPK3 deficiency or catalytically inactive RIPK1 provides greater benefit than MLKL deficiency in mouse models of inflammation and tissue injury. Cell Death Differ. 23:1565–1576. doi:10.1038/cdd.2016.46.

Nguyen, T., and J. Russell. 2001. The regulation of FasL expression during activation-induced cell death (AICD). Immunology. 103:426–434. doi:10.1046/j.1365-2567.2001.01264.x.

Orozco, S., N. Yatim, M.R. Werner, H. Tran, S.Y. Gunja, S.W.G. Tait, M.L. Albert, D.R. Green, and A. Oberst. 2014. RIPK1 both positively and negatively regulates RIPK3 oligomerization and necroptosis. Cell Death Differ. 21:1511–1521. doi:10.1038/cdd.2014.76.

Protasio Veras, F., M. Cornejo Pontelli, C. Meirelles Silva, J.E. Toller-Kawahisa, M. de Lima, D. Carvalho Nascimento, A. Henriques Schneider, D. Caetite, L. Alves Tavares, I.M. Paiva, R. Rosales, D. Colon, R. Martins, I. Araujo Castro, G.M. Almeida, M.I. Fernandes Lopes, M. Nilson Benatti, L. Pastorelli Bonjorno, M. Cavichioli Giannini, R. Luppino-Assad, S. Luna Almeida, F. Vilar, R. Santana, V.R. Bollela, M. Auxiliadora-Martins, M. Borges, C. Henrique Miranda, A. Pazin-Filho, L. Lamberti P. da Silva, L. Dias Cunha, D.S. Zamboni, F. Dal-Pizzol, L.O. Leiria, L. Siyuan, S. Batah, A. Fabro, T. Mauad, M. Dolhnikoff, A. Duarte-Neto, P. Saldiva, T. Mattar Cunha, J.C. Alves-Filho, E. Arruda, P. Louzada-Junior, R. Donizeti Oliveira, and F. Queiroz Cunha. 2020. SARS-CoV-2 triggered neutrophil extracellular traps (NETs) mediate COVID-19 pathology. J Exp Med. 217:e20201129. doi:10.1101/2020.06.08.20125823.

Qin, C., L. Zhou, Z. Hu, S. Zhang, S. Yang, Y. Tao, C. Xie, K. Ma, K. Shang, W. Wang, and D.S. Tian. 2020. Dysregulation of immune response in patients with coronavirus 2019 (COVID-19) in Wuhan, China. Clin. Infect. Dis. 71:762–768. doi:10.1093/cid/ciaa248.

van Raam, B.J., A. Drewniak, V. Groenewold, T.K. Van Den Berg, and T.W. Kuijpers. 2008. Granulocyte colony-stimulating factor delays neutrophil apoptosis by inhibition of calpains upstream of caspase-3. Blood. 112:2046–2054. doi:10.1182/blood-2008-04-149575.

Radermecker, C., N. Detrembleur, J. Guoit, E. Cavalier, M. Henket, C. D’Emal, C. Vanwinge, D. Cataldo, C. Oury, P. Delvenne, and T. Marichall. 2020. Neutrophil extracellular traps infiltrate the lung airway, interstitial and vascular compartments in severe Covid-19. J Exp Med. 217:e20201012s.

Rodrigues, T.S., K.S.G. de Sa, A.Y. Ishimoto, A. Becerra, S. Oliveira, L. Almeida, A. V. Goncalves, D.B. Perucello, W.A. Andrade, R. Castro, F.P. Veras, J.E. Toller-Kawahisa, D.C. Nascimento, M.H.F. de Lima, C.M.S. Silva, D.B. Caetite, R.B. Martins, I.A. Castro, M.C. Pontelli, F.C. de Barros, N.B. do Amaral, M.C. Giannini, L.P. Bonjorno, M.I.F. Lopes, R.C. Santana, F.C. Vilar, M. Auxiliadora-Martins, R. Luppino-Assad, S.C.L. de Almeida, F.R. de Oliveira, S.S. Batah, L. Siyuan, M.N. Benatti, T.M. Cunha, J.C. Alves-Filho, F.Q. Cunha, L.D. Cunha, F.G. Frantz, T. Kohlsdorf, A.T. Fabro, E. Arruda, de O.R.D. R, P. Louzada-Junior, and D.S. Zamboni. 2020. Inflammasomes are activated in response to SARS-CoV-2 infection and are associated with COVID-19 severity in patients. J Exp Med. 218:e20201707.

Ronchetti, S., E. Ricci, G. Migliorati, M. Gentili, and C. Riccardi. 2018. How glucocorticoids affect the neutrophil life. Int. J. Mol. Sci. 19. doi:10.3390/ijms19124090.

Schilling, R., P. Geserick, and M. Leverkus. 2014. Characterization of the ripoptosome and its components: Implications for anti-inflammatory and cancer therapy. 2014. Methods Enzymol. 545:83–102. doi: 10.1016/B978-0-12-801430-1.00004-4.

Schneider, C.A., W.S. Rasband, and K.W. Eliceiri. 2012. NIH Image to ImageJ: 25 years of image analysis. Nat. Methods. 9:671–675. doi:10.1038/nmeth.2089.

Schulte-Schrepping, J., N. Reusch, D. Paclik, K. Baßler, S. Schlickeiser, B. Zhang, B. Krämer, T. Krammer, S. Brumhard, L. Bonaguro, E. De Domenico, D. Wendisch, M. Grasshoff, T.S. Kapellos, M. Beckstette, T. Pecht, A. Saglam, O. Dietrich, H.E. Mei, A.R. Schulz, C. Conrad, D. Kunkel, E. Vafadarnejad, C.J. Xu, A. Horne, M. Herbert, A. Drews, C. Thibeault, M. Pfeiffer, S. Hippenstiel, A. Hocke, H. Müller-Redetzky, K.M. Heim, F. Machleidt, A. Uhrig, L. Bosquillon de Jarcy, L. Jürgens, M. Stegemann, C.R. Glösenkamp, H.D. Volk, C. Goffinet, M. Landthaler, E. Wyler, P. Georg, M. Schneider, C. Dang-Heine, N. Neuwinger, K. Kappert, R. Tauber, V. Corman, J. Raabe, K.M. Kaiser, M.T. Vinh, G. Rieke, C. Meisel, T. Ulas, M. Becker, R. Geffers, M. Witzenrath, C. Drosten, N. Suttorp, C. von Kalle, F. Kurth, K. Händler, J.L. Schultze, A.C. Aschenbrenner, Y. Li, J. Nattermann, B. Sawitzki, A.E. Saliba, L.E. Sander, A. Angelov, R. Bals, A. Bartholomä us, A. Becker, D. Bezdan, E. Bonifacio, P. Bork, T. Clavel, M. Colome-Tatche, A. Diefenbach, A. Dilthey, N. Fischer, K. Förstner, J.S. Frick, J. Gagneur, A. Goesmann, T. Hain, M. Hummel, S. Janssen, J. Kalinowski, R. Kallies, B. Kehr, A. Keller, S. Kim-Hellmuth, C. Klein, O. Kohlbacher, J.O. Korbel, et al. 2020. Severe COVID-19 Is Marked by a Dysregulated Myeloid Cell Compartment. Cell. 182:1419-1440.e23. doi:10.1016/j.cell.2020.08.001.

Schultheiß, C., L. Paschold, D. Simnica, M. Mohme, E. Willscher, L. von Wenserski, R. Scholz, I. Wieters, C. Dahlke, E. Tolosa, D.G. Sedding, S. Ciesek, M. Addo, and M. Binder. 2020. Next-Generation Sequencing of T and B Cell Receptor Repertoires from COVID-19 Patients Showed Signatures Associated with Severity of Disease. Immunity. 53:442-455.e4. doi:10.1016/j.immuni.2020.06.024.

Schwartz, J.T., J.H. Barker, J. Kaufman, D.C. Fayram, J.M. McCracken, and L.-A.H. Allen. 2012. Francisella tularensis Inhibits the Intrinsic and Extrinsic Pathways To Delay Constitutive Apoptosis and Prolong Human Neutrophil Lifespan. J. Immunol. 188:3351– 3363. doi:10.4049/jimmunol.1102863.

Serrao, K.L., J.D. Fortenberry, M.L. Owens, F.L. Harris, and L.A.S. Brown. 2001. Neutrophils induce apoptosis of lung epithelial cells via release of soluble Fas ligand. Am. J. Physiol. - Lung Cell. Mol. Physiol. 280:298–305. doi:10.1152/ajplung.2001.280.2.l298.

Suda, T., H. Hashimoto, M. Tanaka, T. Ochi, and S. Nagata. 1997. Membrane Fas ligand kills human peripheral blood T lymphocytes, and soluble fas ligand blocks the killing. J. Exp. Med. 186:2045–2050. doi:10.1084/jem.186.12.2045.

Tummers, B., L. Mari, C.S. Guy, B.L. Heckmann, D.A. Rodriguez, S. Rühl, J. Moretti, J.C. Crawford, P. Fitzgerald, T.-D. Kanneganti, L.J. Janke, S. Pelletier, J.M. Blander, and D.R. Green. 2020. Caspase-8-Dependent Inflammatory Responses Are Controlled by Its Adaptor, FADD, and Necroptosis. Immunity. 52:1–13. doi:10.1016/j.immuni.2020.04.010.

Varga, Z., A.J. Flammer, P. Steiger, M. Haberecker, R. Andermatt, A.S. Zinkernagel, M.R. Mehra, R.A. Schuepbach, F. Ruschitzka, and H. Moch. 2020. Endothelial cell infection and endotheliitis in COVID-19. Lancet. 395:1417–1418. doi:10.1016/S0140-6736(20)30937-5.

Wang, X., Z. He, H. Liu, S. Yousefi, and H.-U. Simon. 2016. Neutrophil Necroptosis Is Triggered by Ligation of Adhesion Molecules following GM-CSF Priming. J. Immunol. 197:4090–4100. doi:10.4049/jimmunol.1600051.

Wang, X., S. Yousefi, and H.U. Simon. 2018. Necroptosis and neutrophil-Associated disorders review-Article. Cell Death Dis. 9. doi:10.1038/s41419-017-0058-8.

Wendel Garcia, P.D., T. Fumeaux, P. Guerci, D.M. Heuberger, J. Montomoli, F. Roche-Campo, R.A. Schuepbach, and M.P. Hilty. 2020. Prognostic factors associated with mortality risk and disease progression in 639 critically ill patients with COVID-19 in Europe: Initial report of the international RISC-19-ICU prospective observational cohort. EClinicalMedicine. 25:1–11. doi:10.1016/j.eclinm.2020.100449.

Wunsch, H. 2020. Mechanical Ventilation in COVID-19: Interpreting the Current Epidemiology. Am. J. Respir. Crit. Care Med. 202:1–4. doi:10.1164/rccm.202004-1385ED.

Yang, J.W., L. Yang, R.G. Luo, and J.F. Xu. 2020. Corticosteroid administration for viral pneumonia: COVID-19 and beyond. Clin. Microbiol. Infect. 26:1171–1177. doi:10.1016/j.cmi.2020.06.020.

Yipp, B.G., B. Petri, D. Salina, C.N. Jenne, B.N.V. Scott, L.D. Zbytnuik, K. Pittman, M. Asaduzzaman, K. Wu, H.C. Meijndert, S.E. Malawista, A. De Boisfleury Chevance, K. Zhang, J. Conly, and P. Kubes. 2012. Infection-induced NETosis is a dynamic process involving neutrophil multitasking in vivo. Nat. Med. 18:1386–1393. doi:10.1038/nm.2847.

Zhou, Y., C. Niu, B. Ma, X. Xue, Z. Li, Z. Chen, F. Li, S. Zhou, X. Luo, and Z. Hou. 2018. Inhibiting PSMα-induced neutrophil necroptosis protects mice with MRSA pneumonia by blocking the agr system. Cell Death Dis. 9. doi:10.1038/s41419-018-0398-z.

Zhu, L., L. Liu, Y. Zhang, L. Pu, J. Liu, X. Li, Z. Chen, Y. Hao, B. Wang, J. Han, G. Li, S. Liang, H. Xiong, H. Zheng, A. Li, J. Xu, and H. Zeng. 2018. High Level of Neutrophil Extracellular Traps Correlates with Poor Prognosis of Severe Influenza A Infection. J. Infect. Dis. 217:428–437. doi:10.1093/infdis/jix475.

Zhu, L., P. Yang, Y. Zhao, Z. Zhuang, Z. Wang, R. Song, J. Zhang, C. Liu, Q. Gao, Q. Xu, X. Wei, H.X. Sun, B. Ye, Y. Wu, N. Zhang, G. Lei, L. Yu, J. Yan, G. Diao, F. Meng, C. Bai, P. Mao, Y. Yu, M. Wang, Y. Yuan, Q. Deng, Z. Li, Y. Huang, G. Hu, Y. Liu, X. Wang, Z. Xu, P. Liu, Y. Bi, Y. Shi, S. Zhang, Z. Chen, J. Wang, X. Xu, G. Wu, F.S. Wang, G.F. Gao, L. Liu, and W.J. Liu. 2020. Single-Cell Sequencing of Peripheral Mononuclear Cells Reveals Distinct Immune Response Landscapes of COVID-19 and Influenza Patients. Immunity. 53:685-696.e3. doi:10.1016/j.immuni.2020.07.009.

