## Supplemental material for "Blunted Fas signaling favors RIPK1-driven neutrophil necroptosis in critically ill COVID-19 patients"

**Table S1. COVID-19 patients' demographics and clinical characteristics.**

| **Demographics** | | **%** |
|---|---|---|
| Patients (n) | 61 | |
| Age, median ± SD (years) | 64 ± 11.5 | |
| Female | 17 | 28% |
| **Comorbidities** | | |
| BMI[a], median ± SD (kg/m$^2$) | 28 ± 6.2 | |
| CCI[b], median ± SD | 1 ± 1.6 | |
| Diabetes | 23 | 38% |
| COPD, Asthma | 9 | 15% |
| Malignancy | 5 | 8% |
| Immunosuppression | 8 | 13% |
| **Medications** | | |
| Steroids (any) | 48 | 79% |
| *Dexamethasone* | *43* | *70%* |
| Hydroxychloroquine | 7 | 11% |
| Remdesivir | 26 | 43% |
| Antibodies | 2 | 3% |
| Antibiotics | 57 | 93% |
| **Laboratorial findings,** median ± SD | | |
| CRP (mg/l)[c] | 92.5 ± 106.5 | |
| LDH (U/l)[d] | 708.5 ± 287 | |
| IL-6 (ng/l) | 31.7 ± 196.8 | |
| D-Dimer (µg/ml)[e] | 2.1 ± 3.5 | |
| Ferritin (ng/ml)[f] | 1149 ± 1432.3 | |
| Fibrinogen (g/l)[g] | 5.7 ± 1.2 | |
| Myoglobin (ng/ml)[h] | 86 ± 140.5 | |
| Bilirubin (µmol/l)[i] | 6 ± 7.4 | |
| Haemoglobin (G/l)[j] | 115 ± 17.9 | |
| Leucocytes (G/l)[k] | 10.2 ± 6.5 | |
| Lymphocytes (G/l)[l] | 0.9 ± 1.2 | |
| Neutrophils (G/l)[m] | 7.9 ± 4.9 | |
| Monocytes (G/l)[n] | 0.6 ± 0.5 | |
| Platelets (G/ml)[o] | 277.5 ± 113.1 | |
| **Respiratory status** | | |
| Mechanical ventilation | 56 | 92% |
| **Outcome** | | |
| 30 day survival | 52 | 85% |
| ICU discharge survival | 51 | 84% |

*[a]BMI: Body mass index; [b]CCI: Charlson comorbidity index; [c]CRP: C-reactive protein (normal value <5 mg/l); [d]LDH: lactate dehydrogenase (normal range: 240–480 U/l); [e]D-dimers (normal value <0.5 µg/ml); [f]Ferritin (normal range: 30-400 ng/ml); [g]Fibrinogen (normal range: 2-4 g/l); [h]Myoglobin (normal range: 28-72 ng/ml); [i]Bilirubin (normal value < 2 1µmol/l); [j]Haemoglobin (normal range: 120-175 g/l); [k]Leucocytes (normal range: 4.00-11.0 G/l); [l]Lymphocytes (normal range: 1.5–3.5 G/l); [m]Neutrophils (normal range: 2.5–7.5 G/l); [n]Monocytes (normal range: 0.30-0.90 G/l); [o]Platelets (normal range: 150-400 G/l)*

**Table S2. Background information of obtained biopsies.**

| |
|---|
| **Thrombus – COVID-19 Patient #1** |
| The fresh thrombus biopsy was obtained from a critically ill 65 years old female COVID-19 patient. The thrombus was located in the groin region. |
| **Thrombus – COVID-19 Patient #2** |
| The thrombus biopsy was obtained from a critically ill 58 years old male COVID-19 patient. The thrombus was located in the A. poplitea of the right limb. |
| **Thrombus – Control (non-COVID-19) Patient** |
| The control thrombus was obtained from a 82 years old female non-COVID-19 patient who suffered from an aneurysm in the A. poplitea of the right limb. |
| **Lung biopsy – COVID-19 Patient #3** |
| The lung biopsy was obtained from a critically ill 59 years old male COVID-19 patient with a pneumothorax following bilateral COVID-19 pneumonia. Of note, the patient furthermore suffered a secondary pulmonary *Pseudomonas aeruginosa* infection, which had resolved at the time of the biopsy. |

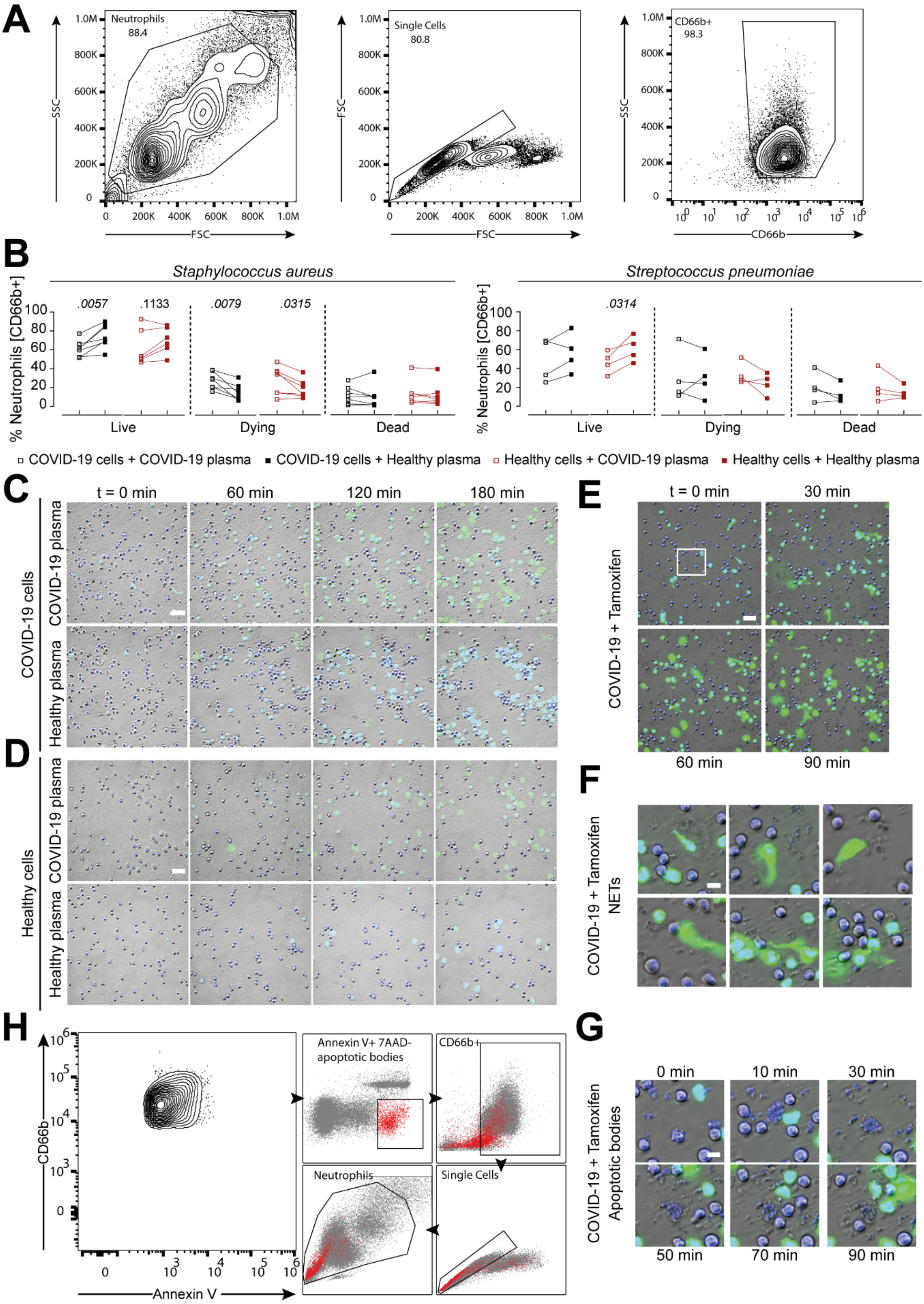
A
Neutrophils
88.4
Single Cells
80.8
CD66b+
98.3
SSC
FSC
CD66b
B
Staphylococcus aureus
Streptococcus pneumoniae
% Neutrophils [CD66b+]
.0057
.1133
.0079
.0315
.0314
Live
Dying
Dead
COVID-19 cells + COVID-19 plasma
COVID-19 cells + Healthy plasma
Healthy cells + COVID-19 plasma
Healthy cells + Healthy plasma
C
t = 0 min
60 min
120 min
180 min
COVID-19 cells
COVID-19 plasma
Healthy plasma
D
Healthy cells
COVID-19 plasma
Healthy plasma
E
t = 0 min
30 min
60 min
90 min
COVID-19 + Tamoxifen
F
COVID-19 + Tamoxifen
NETs
G
0 min
10 min
30 min
50 min
70 min
90 min
COVID-19 + Tamoxifen
Apoptotic bodies
H
CD66b
Annexin V
Annexin V+ 7AAD-
apoptotic bodies
CD66b+
Neutrophils
Single Cells

**Figure S1.**

**(A)** Neutrophil gating strategy. **(B)** Quantification of live (Annexin V-/7AAD-), dying (Annexin V+/7AAD-) and dead (Annexin V+/7AAD+) COVID-19 (n=4-7) and healthy donor neutrophils (n=4-6) pre-stimulated with auto- or heterologous plasma for 2.5 h and subsequently challenged with *S. aureus* or *S. pneumoniae* for 1.5 h. **(C)** Overview of *r*epresentative zoomed in time lapse microscopy of acute COVID-19 neutrophils stimulated with their own or healthy donor plasma as shown in Fig. 1E. Cells were stained with Hoechst 33342 and SYTOX$^{TM}$ green. Images were taken every ten minutes. Scale bar indicates 30 µm. **(D)** Overview of representative time lapse microscopy of healthy neutrophils stimulated with auto- or heterologous plasma. Scale bar indicates 30µm. (**E**) Time lapse microscopy of a COVID-19 patient which was on additional Tamoxifen treatment. Scale bar indicates 30 µm. **(F)** Images of NETs taken at timepoint 30 min. Scale bar indicates 15 µm. **(G)** Enlarged insert from **(E)**, showing active apoptosis ongoing. Scale bar indicates 10 µm. **(H)** Flow cytometry backgating plot, confirming CD66b+ apoptotic bodies observed in the COVID-19 Tamoxifen patient. Connected squares represent one donor. Statistics were calculated by paired t-test or Wilcoxon signed-rank test. P values are indicated within the graphs.

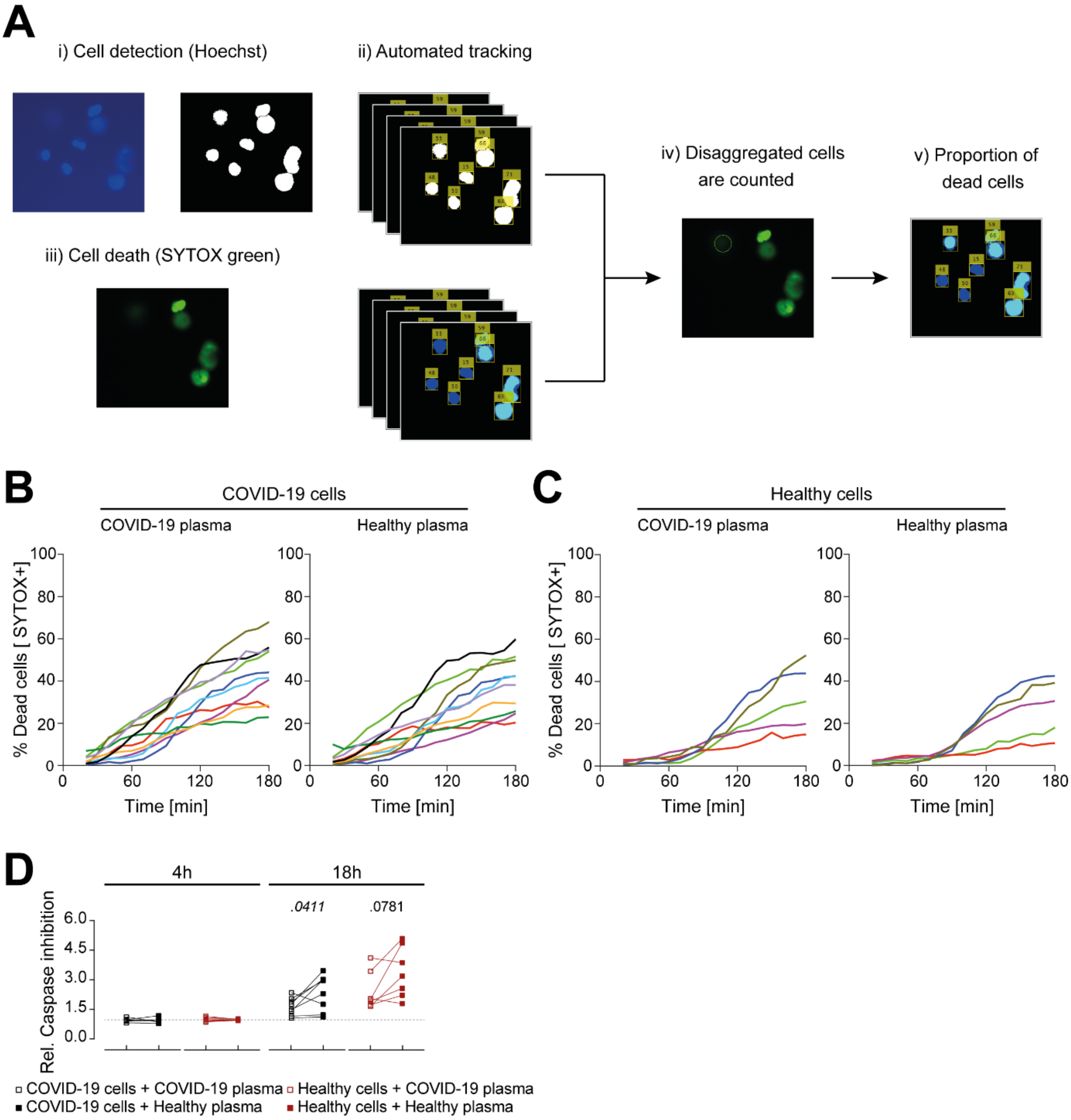
A
i) Cell detection (Hoechst)
ii) Automated tracking
iii) Cell death (SYTOX green)
iv) Disaggregated cells are counted
v) Proportion of dead cells
B
COVID-19 cells
COVID-19 plasma
Healthy plasma
% Dead cells [ SYTOX+]
Time [min]
C
Healthy cells
COVID-19 plasma
Healthy plasma
% Dead cells [ SYTOX+]
Time [min]
D
4h
18h
.0411
.0781
Rel. Caspase inhibition
COVID-19 cells + COVID-19 plasma
COVID-19 cells + Healthy plasma
Healthy cells + COVID-19 plasma
Healthy cells + Healthy plasma

**Figure S2.**

**(A)** Image analysis process used to quantify the microscopy time lapses. Neutrophils were discovered on the Hoechst channel through thresholding (i). Neutrophil tracking was done using Kalman filtering (ii), cell death was counted based on SYTOX$^{TM}$ green penetration when the cell membrane lost integrity (iii). Based on tracking, cells that were counted dead were taken into account even when either signal is lost (iv). The proportion of dead cells was then extracted from each image (v). (**B and C**) Cell death curves of (B) COVID-19 (C) and healthy donor neutrophils stimulated with COVID-19 or healthy plasma for 3 h. Each line indicates one experiment (i.e. donor) as mean of 8 FOVs per condition. (**D**) Relative caspase-inhibition effect on survival of COVID-19 (n=6-8) and healthy donor neutrophils (n=6-7) stimulated with auto- or heterologous plasma at 4 h and 18 h post-stimulation. Each dot, square or connected squares represent one donor. Shown are mean ± SEM. Statistics were calculated by paired t-test or Wilcoxon signed-rank test. FOV, field of view.

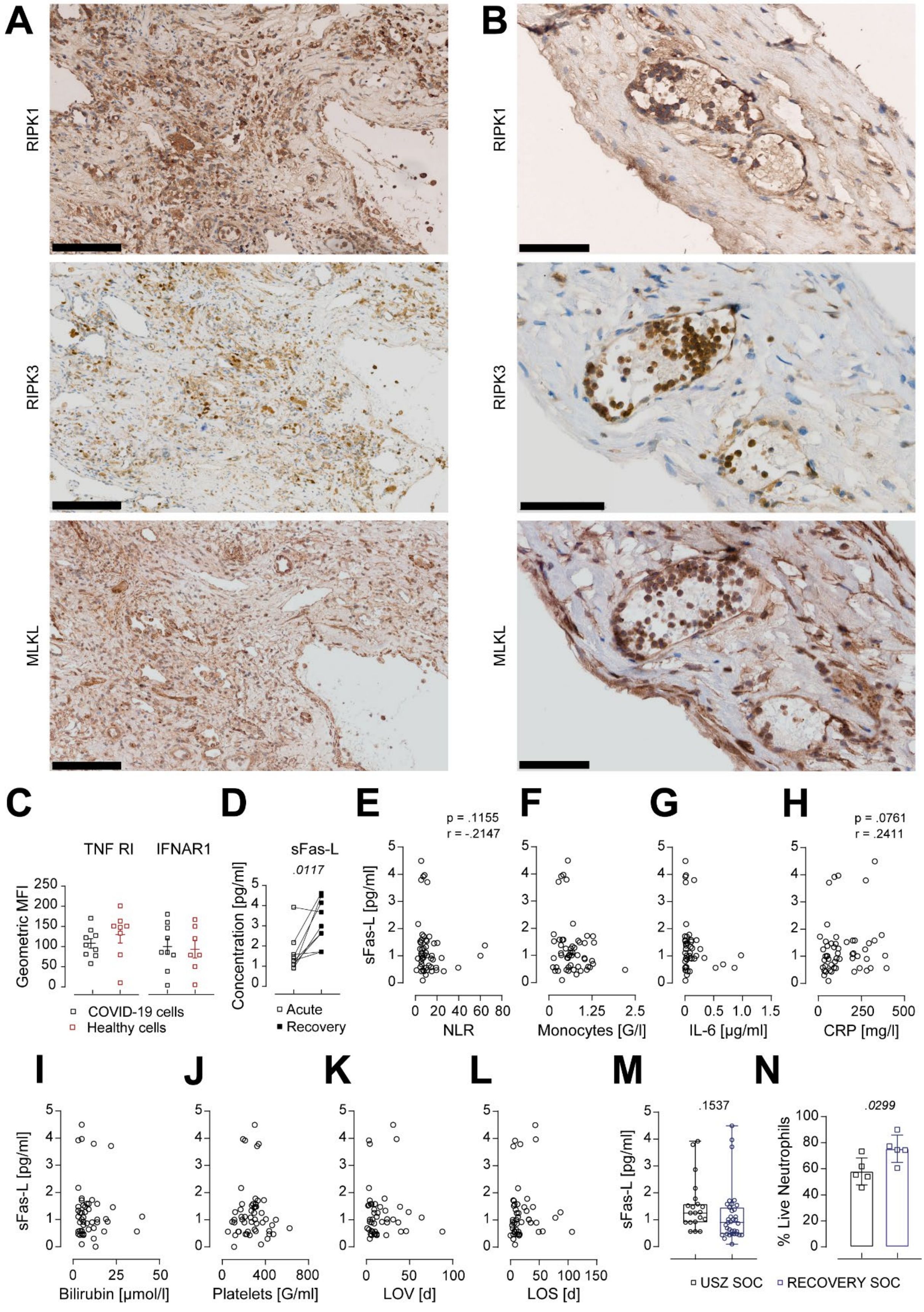
A
B
RIPK1
RIPK3
MLKL
C
TNF RI
IFNAR1
Geometric MFI
COVID-19 cells
Healthy cells
D
sFas-L
.0117
Concentration [pg/ml]
Acute
Recovery
E
p = .1155
r = -.2147
sFas-L [pg/ml]
NLR
F
Monocytes [G/l]
G
IL-6 [µg/ml]
H
p = .0761
r = .2411
CRP [mg/l]
I
Bilirubin [µmol/l]
J
Platelets [G/ml]
K
LOV [d]
L
LOS [d]
M
.1537
N
.0299
% Live Neutrophils
USZ SOC
RECOVERY SOC

**Figure S3.**

(**A-B**) Overview of RIPK1, RIPK3 and MLKL stained COVID-19 lung tissue (A) and neutrophils within blood vessel in the lung (B). Scale bars, 200 µm (A) and 100 µm (B). **(C)** TNF RI and IFNAR1 Receptor expression on COVID-19 (n=5-9) and healthy neutrophils (n=5-8). **(D)** Plasma levels of sFas-L in COVID-19 patients during acute and corresponding recovery phase (n=9). **(E-L)** Correlation analysis of plasma sFas-L levels with NLR (E), monocytes (F), IL-6 (G), CRP (H), bilirubin (I), platelets (J), LOV (K) and LOS (L). (**M**) Plasma sFas-L levels of USZ SOC (n=20) and RECOVERY SOC (n=35). (**N**) In-depth analysis of neutrophil viability from Fig. 1E, according to USZ SOC (n=5) or RECOVERY SOC (n=5). Each dot, square or connected squares represent one donor. Shown are mean ± SEM. Statistics were calculated by paired t-test, Wilcoxon signed-rank test or non-parametric Spearman correlation. P values are indicated within the graphs. SEM, standard error of means; NLR, neutrophil to lymphocyte ratio; CRP, c-reactive protein; LOV, length of ventilation; LOS, length of hospital stay.

**Video 1. Time lapse of COVID-19 neutrophils stimulated with COVID-19 or healthy donor plasma.**

Time lapse microscopy of COVID-19 neutrophils stimulated with auto- or heterologous plasma. Cells were stained with Hoechst 33342 and SYTOX$^{TM}$ green. Images were taken every ten minutes and used for a time lapse. Scale bar, 30 µm. See also Fig.1 and S1.

**Video 2. Time lapse of healthy donor neutrophils stimulated with COVID-19 or healthy donor plasma.**

Time lapse microscopy of healthy donor neutrophils stimulated with auto- or heterologous plasma. Cells were stained with Hoechst 33342 and SYTOX$^{TM}$ green. Images were taken every ten minutes and used for a time lapse. Scale bar, 30 µm. See also Fig. S1.
